# BIG participates in the Arg/N-degron pathways and the hypoxia response in *Arabidopsis thaliana*

**DOI:** 10.1101/2023.05.26.542459

**Authors:** Hongtao Zhang, Chelsea Rundle, Nikola Winter, Alexandra Miricescu, Brian C. Mooney, Andreas Bachmair, Emmanuelle Graciet, Frederica L. Theodoulou

## Abstract

BIG (also known as DOC1 and TIR3) is an 0.5 MDa protein that has been associated with multiple important functions in signalling and development through forward genetic screens in *Arabidopsis thaliana*. However, the biochemical function(s) of BIG are unknown. Here, we investigated whether BIG plays a role in the Arg/N-degron pathways, protein regulatory mechanisms in which substrate protein fate is influenced by the N-terminal (Nt) residue. In Arabidopsis, PROTEOLYSIS1 (PRT1) is an E3 ligase with specificity for aromatic amino acids, whereas PROTEOLYSIS6 (PRT6) targets basic N-terminal residues. We crossed a big loss-of-function allele to *prt6* and *prt1* mutants and examined the stability of protein substrates. Stability of model N-degron pathway substrates was enhanced in *prt6-1 big-2* and *prt1-1 big-2* relative to the respective single mutants. Abundance of the PRT6 physiological substrates, HYPOXIA RESPONSIVE ERF (HRE)2 and VERNALIZATION (VRN)2 was similarly increased in *prt6 big* double mutants, without increase in transcripts. Accordingly, hypoxia marker expression was enhanced in *prt6 big* double mutants, in a manner requiring arginyltransferase activity and RAP-type ERFVII transcription factors. Transcriptomic analysis of roots not only demonstrated synergistically increased expression of a plethora of hypoxia responsive genes in the double mutant relative to *prt6* but also revealed other roles for PRT6 and BIG, including regulation of suberin deposition through both ERFVII-dependent and independent mechanisms, respectively. Our results show that BIG acts together with PRT6 to regulate the hypoxia response and wider processes.

**Significance Statement:** The N-degron pathways are a group of protein regulatory mechanisms that play important roles in plant growth, development, and response to biotic and abiotic stresses. Despite rapid progress in the last decade, key enzymatic components of the pathways remain to be identified. BIG (also known as DOC1 and TIR3) is a protein of approximately 0.5 MDa, associated with multiple, distinct roles in plants but the precise biochemical functions of this protein have remained enigmatic until now. Here we identify BIG as a new component of plant N-degron pathways that acts together with the N-recognin E3 ligase PROTEOLYSIS6 (PRT6) to control the hypoxia response and other functions in *Arabidopsis thaliana*.

## Introduction

Targeted protein degradation is an important proteostatic mechanism that influences a multitude of agronomically important traits in plants [1,2] and represents a major target for drug development in humans [3,4]. The Arg/N-degron pathways (formerly known as the Arg/N-end rule pathway) constitute a specialised form of proteostasis in which the N-terminal (Nt) residue of a given protein is the key determinant of a degradation signal, known as an N-degron [5,6]. N-degrons are revealed by protein cleavage by non-processive endopeptidases and/or created by subsequent enzymatic modification of the neo-N-terminus by amidases and arginyl tRNA transferase enzymes (ATEs) (Fig. 1A). In mammals and yeast, N-degrons include type 1 positively charged residues (Arg, Lys, His) and type 2 bulky hydrophobic residues (Trp, Phe, Tyr, Leu, and Ile), which are recognised by proteins known as N-recognins that facilitate substrate degradation. The prototypical N-recognin, yeast ubiquitin amino-end recognising protein 1 (UBR1) accepts both type 1 and type 2 substrates, whereas in mammals, four N-recognins, UBR1, UBR2, UBR4/p600 and UBR5, which share a conserved UBR box domain, act semi-redundantly to mediate proteasomal and autophagic degradation of Arg/N-degron pathway substrates [7-9]. In contrast, plants contain N-recognins with discrete substrate specificities [10]. PRT6, the Arabidopsis homologue of ScUBR1 and HsUBR1/2, is an E3 ligase with specificity for basic N-termini (Arg, Lys, His), PRT1 is an unrelated ZZ domain protein with specificity for aromatic N-termini (Phe, Tyr, Trp), and experimental evidence indicates the existence of a further (still unknown) N-recognin class that targets bulky/hydrophobic-N-termini (Leu and Ile) [10-13].

**Figure 1.**
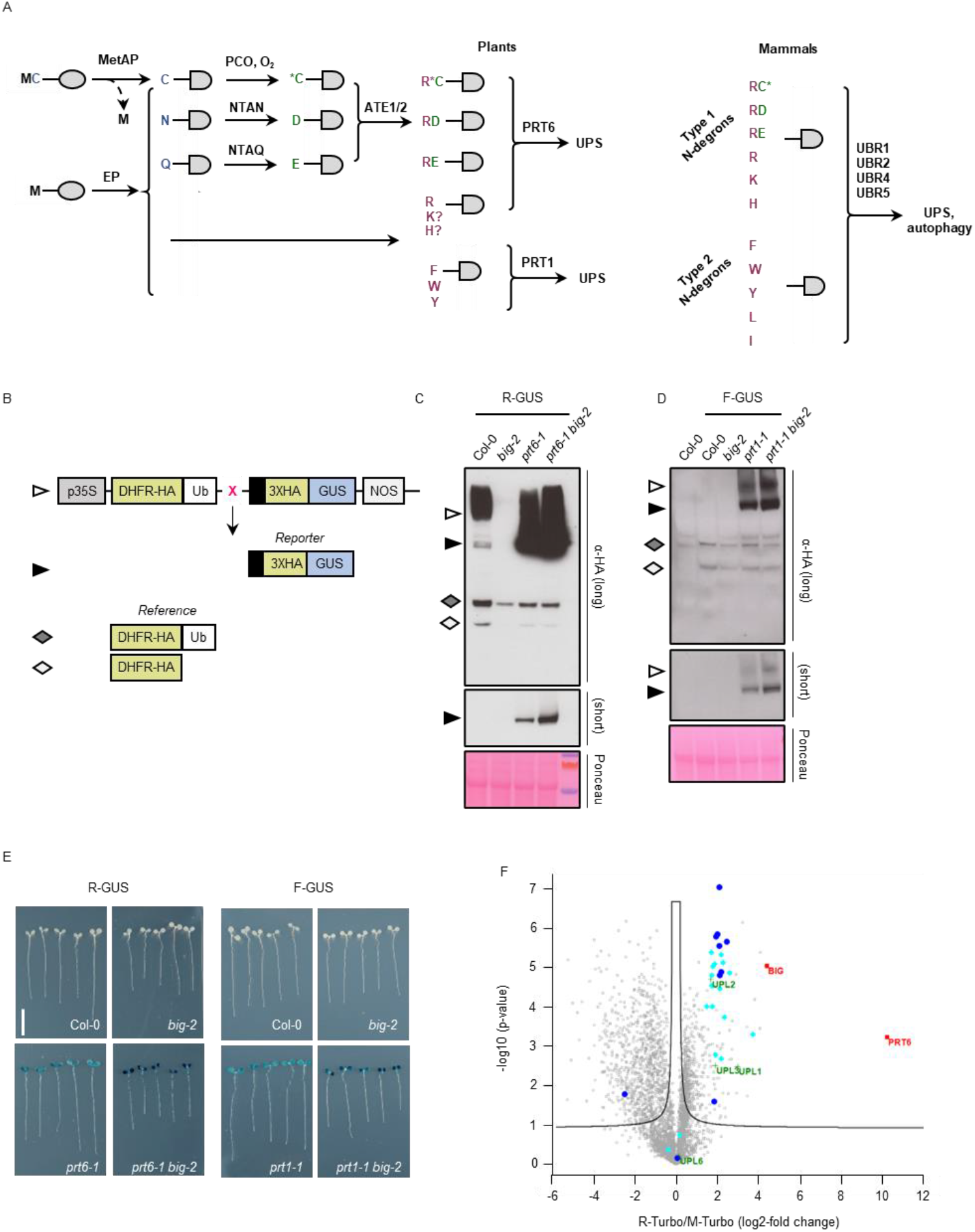
BIG influences the stability of model type 1 and type 2 Arg/N-degron pathway substrates. (A) Schematic showing the architecture of the Arg/N-degron pathway and specificity of N-recognins in plants and mammals. Single letter codes for amino acid residues are used; C* indicates oxidised cysteine. Proteins (represented by grey ovals) may become N-degron pathway substrates via cleavage by non-processive endopeptidases (EP), or by methionine aminopeptidase (MetAP), where the second residue is small. Substrates may also be generated by enzymatic modification of N-termini by PLANT CYSTEINE OXIDASE (PCO), Asn-specific N-terminal amidase (NTAN), Gln-specific N-terminal amidase (NTAQ), and arginyl-tRNA protein transferase (ATE). In plants, destabilising residues thus generated are targeted for degradation by the Ubiquitin Proteasome System (UPS) via N-recognin E3 ligases PROTEOLYSIS6 (PRT6; specific for basic N-termini) and PROTEOLYSIS1 (PRT1; specific for aromatic N-termini). In mammals, four N-recognins act semi-redundantly to mediate the degradation of both type 1 and type 2 substrates via the UPS or by autophagy. (B) Generation of N-degron pathway substrates. Constructs driven by the constitutive *CaMV35S* promoter (p35S) encode a fusion of mouse dihydrofolate reductase (DHFR, yellow) to ubiquitin (variant K48R; Ub, white), followed by *Escherichia coli* beta glucuronidase (GUS, blue). Ubiquitin-specific proteases remove ubiquitin co-translationally to release the GUS reporter protein and reveal a new N terminus (a variable residue, indicated by X). The GUS ORF is extended by unstructured amino acids (black) to enhance the effect of destabilising amino-terminal residues. The cleavage also creates a stable DHFR reference protein and HA epitopes enable immunological detection of both products [10]. NOS, nopaline synthase terminator. (C) Detection of N-degron pathway substrates by immunoblotting of crude protein extracts from 6-d old seedlings of different genotypes expressing R-GUS reporters. Symbols to the left indicate the protein products shown in (B). Blots were developed until the stable reference protein could be detected (α-HA long); where the stabilised reporter band signal is saturated, a shorter exposure is shown in the lower panel (short) for clarity. Ponceau S staining was used to confirm equal loading. (D) As for (C) with F-GUS. (E) Histochemical staining of GUS reporter activity in 6-d old seedlings expressing R-GUS and F-GUS test substrates. Seedlings were rearranged on an agar plate, prior to photography. Bar, 1 cm. (F) Volcano plot visualising enrichment of biotinylated proteins when comparing transgenic lines expressing R-Turbo versus M-Turbo, after proteasome inhibition to ensure equal presence of either protein. PRT6 and BIG are the two most enriched proteins. Turquoise diamonds, non-ATPase subunits of the proteasome; blue circles, ATPase subunits; green crosses, HECT E3 ubiquitin ligases.

The availability of mutants and transgenics in which N-recognin function is disrupted has revealed diverse functions for the Arg/N-degron pathway in plants [14]. Relatively little is known regarding the PRT1/N-degron pathway, although it has been shown to influence immune responses [15,16] and the turnover of the E3 ligase BIG BROTHER [17]. In contrast, the PRT6/N-degron pathway plays multiple roles in development [19-28], plant–pathogen interactions [15,29,30], and responses to the abiotic environment [23, 31-39]. The first substrates of the PRT6/N-degron pathway were identified in the context of oxygen sensing [31,32]. Arabidopsis has five Group VII ethylene response factor transcription factors (ERFVIIs) bearing a conserved cysteine residue at position two, of which *RELATED TO APETALA (RAP) 2*.*12, RAP2*.*2*, and *RAP2*.*3* are constitutively expressed, whereas *HYPOXIA RESPONSIVE ERF* (*HRE*) *1* and *HRE2* are induced by low oxygen [40]. All five Met-Cys-ERFVII proteins undergo co-translational Nt Met excision to reveal Nt Cys, which under normoxia is susceptible to oxidation by PLANT CYSTEINE OXIDASE (PCO) enzymes and subsequent arginylation by ATEs [33,41]. N-terminally arginylated ERFVIIs are then recognised by PRT6, which targets the proteins for proteasomal degradation [31,32]. However, when oxygen is limiting, ERFVIIs are stabilised and co-ordinate the transcriptional response to hypoxia. Consequently, hypoxia responsive genes, such as *ALCOHOL DEHYDROGENASE* (*ADH*) and *PHYTOGLOBIN1* (*PGB1*) (as well as *HRE1* and *HRE2*) are ectopically expressed in *prt6* alleles [19,31,42]. The Arabidopsis genome encodes 248 Met-Cys initiating proteins, of which the polycomb repressive complex 2 subunit, VERNALISATION 2 (VRN2) and the transcription factor LITTLE ZIPPER 2 (ZPR2) have also been confirmed as oxygen-sensitive physiological PRT6/N-degron pathway substrates with roles in development [26,27].

Experimental evidence and sequence database searches indicate that the full suite of N-recognins has not yet been identified in plants [10,13]. Here, we investigated a potential role for the UBR box protein, BIG/TIR3/DOC1 in Arg/N-degron pathways, since its mammalian and Drosophila homologues (UBR4/p600 and Calossin/Pushover, respectively) are known N-recognins [7,8, 43-45]. *BIG* has been identified in around twenty different forward genetic screens and associated with diverse physiological functions via reverse genetics in Arabidopsis. The first *big* allele, *dark overexpression of CAB* (*doc1-1*), was isolated in a screen for mutants with mis-regulated photosynthetic gene expression [46]. *doc1-1*, which displays a striking morphological phenotype of reduced apical dominance and small stature, was subsequently found to be allelic to *transport inhibitor response3* (*tir3-1*), a mutant compromised in auxin transport [47,48]. The affected gene was identified via map-based cloning and renamed in recognition of its exceptional size: *BIG*, which is expressed throughout the plant, encodes a 5077 amino acid protein with a predicted molecular weight of 565,597 Da [48,49]. *BIG* was later shown to influence multiple hormone signalling pathways and different aspects of plant development [50-58]. Recent studies indicate further, apparently disparate functions for *BIG* in the circadian clock, guard cell signalling, calcium homeostasis, regulation of C/N balance, response to pathogens, cell death, and wound-induced rooting [49, 56, 59-64]. Although many *big* mutant phenotypes can be ascribed to dysregulation of auxin transport [46, 47, 48, 52, 53, 54, 57, 58, 65, 66, 67, 68], this is not the case for all processes influenced by BIG and to date its precise biochemical functions have remained unclear.

In this study, we demonstrate that BIG participates in the Arg/N-degron pathways, acting semi-redundantly with PRT6 and PRT1. PRT6/N-degron pathway substrates are hyperaccumulated in *prt6 big* double mutants, enhancing the molecular response to hypoxia in an ERFVII-dependent fashion. This was confirmed by RNA-seq analysis which also indicated a range of different genetic interactions between *big-2* and *prt6-5* that influence transcription of additional groups of genes, pointing to new functions for BIG and PRT6, including the regulation of suberin deposition.

## Results

### BIG influences the stability of model Arg/N-degron pathway substrates

To test whether BIG plays a role in the Arg/N-degron pathways, we used the ubiquitin fusion technique to produce pathway substrates *in planta* [69]. We took advantage of the DHFR-Ub-X-GUS system [10], in which a genetically encoded ubiquitin domain is cleaved *in vivo* by deubiquitinating enzymes to produce a reporter protein, β-glucuronidase (GUS) bearing a residue of choice (X) at the N-terminus, and a stable reference protein, dihydrofolate reductase (DHFR) (Fig. 1B). Lines expressing constructs designed to release a type 1 substrate (R-GUS), a type 2 substate (F-GUS) and a stable control (M-GUS) were generated in the wild-type Arabidopsis accession, Columbia-0 (Col-0), and in different mutant backgrounds lacking known N-recognins and *BIG*. The stability of X-GUS was assessed by immunoblotting and histochemical staining. Immunoblotting revealed that the fusion proteins were cleaved as predicted and that R-GUS and F-GUS were unstable in Col-0 wild type seedlings, relative to the DHFR control (Fig. 1C, D; Fig S1A, B). Removal of N-recognin function resulted in stabilisation of R-GUS and F-GUS in *prt6-1* and *prt1-1* mutants, respectively, as previously reported [10, 24]. In contrast, M-GUS was stable in all backgrounds tested (Fig S1B, C). R- and F-GUS reporters were not stabilised in the *big-2* single mutant, but stability of R-GUS was enhanced in the *prt6-1 big-2* double mutant compared to *prt6-1*. Similarly, F-GUS was more stable in *prt1-1 big-2* than in *prt1-1* (Fig. 1C,D).

Histochemical staining for GUS activity was consistent with these results (Fig. 1E). Together, these data indicate that BIG acts synergistically with known N-recognins to mediate the degradation of substrates initiating with R and F. As an independent test, a cleavable R-luciferase (R-LUC) reporter [13, 70] was also introduced into *prt6-5 big-2* (Fig S1D). R-LUC was unstable in Col-0 and *big-2* but detected in both *prt6-5* and *prt6-5 big-2*, with higher luciferase activity in the double mutant, consistent with enhanced stabilisation of the R-LUC protein in *prt6-5 big-2* (Fig S1E).

To complement the genetic approach, potential protein interactions with N-degrons were investigated using proximity labelling [71]. Transgenic lines expressing a modified *Escherichia coli* biotin ligase (TurboID)-YFP fusion designed to reveal either an Nt M- or R-residue (Fig. S1F) were generated in the Col-0 background. Both PRT6 and BIG were enriched in the R-TurboID sample relative to the M-TurboID sample, indicating their proximity to R-TurboID *in planta*, and suggesting a potential physical interaction of Nt Arg residues with PRT6 and BIG (Fig. 1F).

Interestingly, regulatory proteasomal subunits and Homologous to E6AP C-terminus (HECT) ubiquitin E3 ligases known to be associated with the proteasome [72] were also highly enriched in the R-TurboID sample (Fig. 1F; Dataset S1).

### BIG influences the abundance of physiological PRT6/N-degron pathway substrates

To explore whether BIG influences the stability of physiological substrates, we focused on the PRT6/N-degron pathway, for which several targets have been identified. We first tested two representative ERFVII transcription factors by crossing plants expressing HA-tagged HRE2 and RAP2.3 [22,31] to N-degron pathway mutants and *big-2*. The protein abundance of HRE2-HA was increased in *big-2 prt6-5* roots relative to the single *prt6-5* mutant (Fig. 2A). Transgene-specific q-RT-PCR showed that the increased levels of HRE2-HA protein were not driven by an increase in transcript abundance (Fig. 2B). Consistent with the known role of ERFVIIs in the hypoxia response, the enhanced HRE2-HA protein abundance in *prt6-5 big-2* relative to *prt6-5* was accompanied by a much stronger induction of the core hypoxia genes *ALCOHOL DEHYDROGENASE* (*ADH*) and *PHYTOGLOBIN1* (*PGB1*) (Fig. 2A, C). RAP2.3-HA protein was stable in *prt6-5* seedlings, but not detectable in Col-0 and *big-2* (Fig. 2D) and hypoxia markers were strongly enhanced in the *prt6-5* line (Fig. 2D, F). Transgene transcript levels were lower in the *prt6-5* background compared to Col-0, indicating that the increased abundance of RAP2.3-HA is due to post-transcriptional regulation (Fig. 2E). *prt6-5 big-2* double mutant lines expressing RAP2.3-HA exhibited curled leaves and generally stunted growth, and flowering was extremely delayed with only a short primary bolt produced (Fig. S2A). We were unable to recover seeds from these plants; dissection of flowers revealed incompletely elongated stamens that did not mature or release pollen (Fig. S2B). Therefore, we analysed pooled seedlings homozygous for *prt6-5* but segregating for *big-2*. Accordingly, we observed a modest increase in RAP2.3-HA abundance and hypoxia marker expression, despite only one quarter of these seedlings being homozygous for both mutations (Fig. 2G).

**Figure 2.**
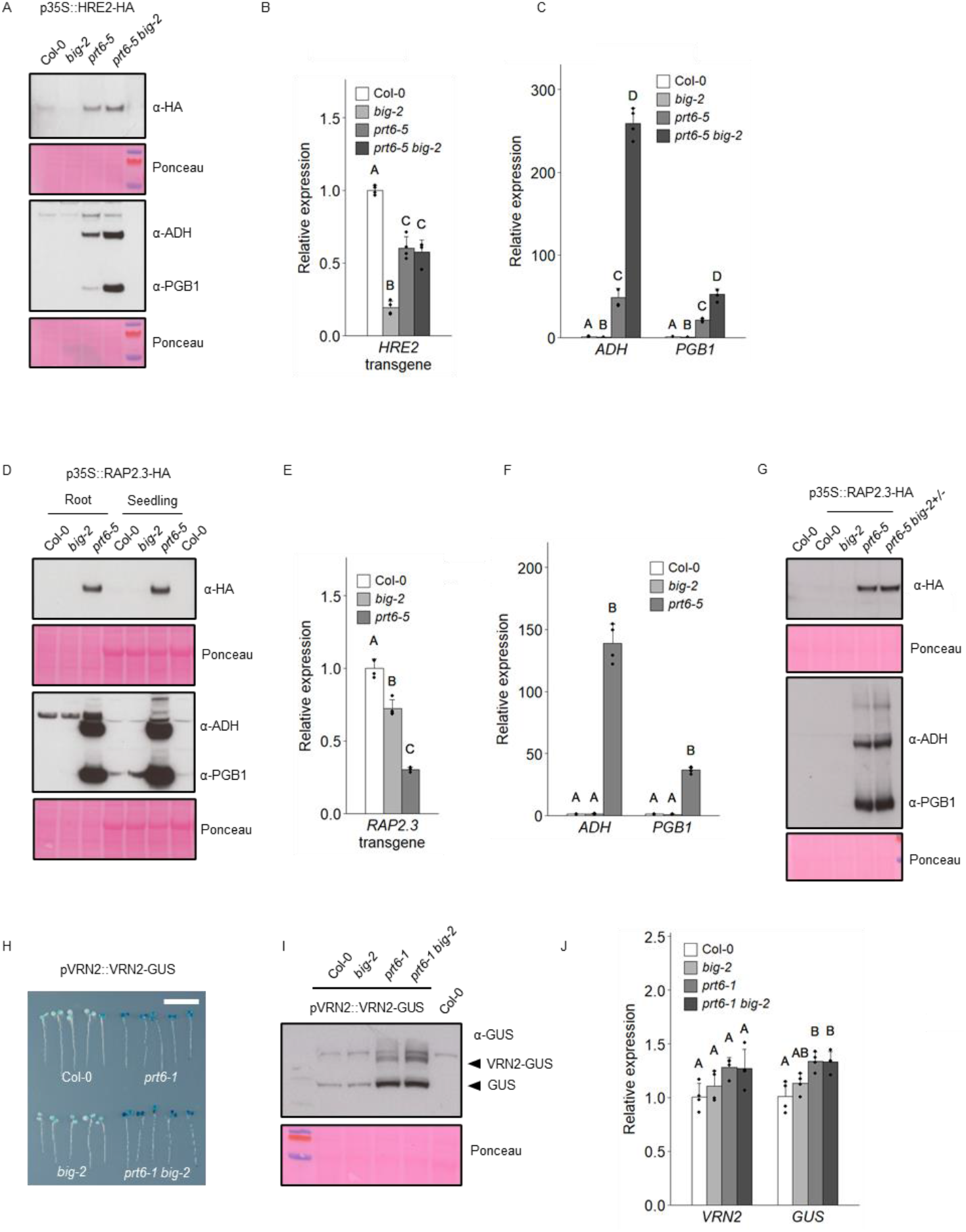
BIG influences the abundance of physiological PRT6/N-degron pathway substrates. (A-C) Molecular analysis of seedlings expressing p35S::HRE2-HA. (A) Immunoblots of crude protein extracts from 6-d old roots of the indicated genotypes, probed with anti-HA (α-HA) antiserum or antibodies specific for the hypoxia markers, ADH and PGB1 (the hypoxia marker antibodies were applied simultaneously to a single membrane). Ponceau S staining was used to confirm equal loading. (B) Relative expression of *HRE2* transgene. Values are means ± SD. (C) Relative expression of *ADH* and *PGB1*; Values are means ± SD. (D-G) Molecular analysis of seedlings expressing p35S::RAP2.3-HA. (D, G) Immunoblots of crude protein extracts from 6-d old seedlings of the indicated genotypes, probed with anti-HA (α-HA) antiserum or antibodies specific for the hypoxia markers, ADH and PGB1. Ponceau S staining was used to confirm equal loading. (E) Relative expression of *RAP2*.*3* transgene. Values are means ± SD. (F) Relative expression of *ADH* and *PGB1*; Values are means ± SD. (H-J) Molecular analysis of seedlings expressing pVRN2::VRN2-GUS. (H) Histochemical staining of GUS reporter activity in 6-d old seedlings. Seedlings were rearranged on an agar plate prior to photography. Bar, 1 cm. (I) Immunoblot of 6-d old seedlings probed with anti-GUS antibody. Ponceau S staining was used to confirm equal loading. (J) Relative expression of *VRN2* and *GUS*. Values are means ± SD. For all plots, different letters indicate significant differences between conditions (*P* < 0.05).

We next tested whether the abundance of a functionally distinct endogenous N-degron pathway substrate, VRN2, was influenced by BIG, using a VRN2-GUS fusion driven by the native *VRN2* promoter [26]. Histochemical staining confirmed previous results [26] with GUS present throughout the seedling in the *prt6-1* background, and additionally revealed increased intensity in *prt6-1 big-2* relative to *prt6-1* (Fig. 2H). Immunoblotting showed specifically that VRN2-GUS was increased in abundance in *prt6-1 big-2* compared to *prt6-1* and unstable in Col-0 and *big-2* (Fig. 2I). q-RT-PCR demonstrated that there were no significant differences in *VRN2 or GUS* expression between *prt6-1* and *prt6-1 big-2*, indicating that changes in VRN2-GUS abundance relate to post-translational control by PRT6 and BIG (Fig. 2J).

### BIG works in parallel with PRT6 to regulate the hypoxia response

To further understand how BIG regulates the hypoxia response, we constructed a series of combination mutants using alleles lacking pathway substrates and enzymes, and quantified hypoxia markers. Firstly, to observe whether arginylation is necessary for BIG to participate in the N-degron pathway, a mutant lacking arginyl transferase activity was crossed to *big-2*. Expression of *ADH* and *PGB1* and accumulation of the respective proteins were comparable in *ate1/2* and *ate1/2 big-2*, indicating that regulation of the hypoxia response by *BIG* is dependent on arginylation (Fig. 3A, B). Genetic removal of *RAP2*.*12, RAP2*.*2* and *RAP2*.*3* was sufficient to prevent constitutive expression of hypoxia markers in *prt6-1* seedlings, as shown previously [24], and also in the *prt6-1 big-2* background (Fig. 3C, D) demonstrating that BIG influences hypoxia gene expression exclusively through RAP-type ERFVIIs. The mutants were also followed through development to determine whether regulation of ERVIIs underpins other phenotypes of *prt6-1 big-2*. Removal of RAP-type ERFVIIs did not have an impact on the overall morphology of *big-2*, consistent with the lack of stabilisation in the single mutant. However, the stunted size and delayed flowering of *prt6-1 big-2* was partially rescued by removal of these substrates (Fig. S3).

**Fig. 3.**
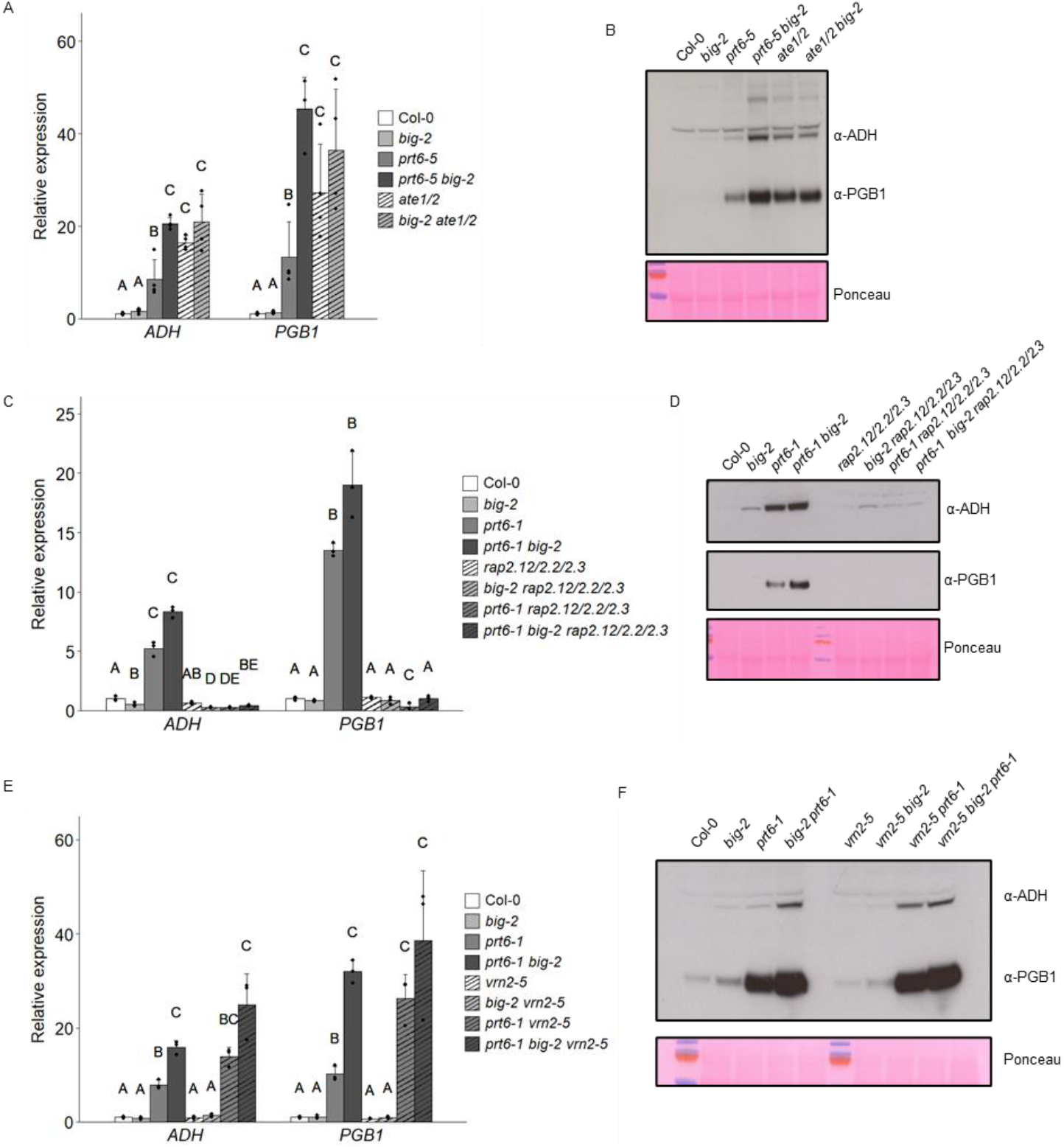
Regulation of the hypoxia response by BIG requires arginylation and ERFVIIs. (A, B) Molecular analysis of PRT6/N-degron pathway mutants. (A) Relative expression of *ADH* and *PGB1* in 6-d old seedlings. Values are means ± SD (n=4). (B) Immunoblots of crude protein extracts from 6-d old seedlings of the indicated genotypes, probed with anti-HA (α-HA) antiserum or antibodies specific for the hypoxia markers, ADH and PGB1 (which were applied to the same membrane). (C-F) Molecular analysis of PRT6/N-degron pathway mutants combined with *rap2*.*12/2*.*2/2*.*3* mutant alleles (C, D) or with *vrn2-5* (E, F). (C, E) Relative expression of *ADH* and *PGB1* in 6-d old seedlings. Values are means± SD (n=3). (D, F) Immunoblot of crude protein extracts from 6-d old seedlings of the indicated genotypes, probed with anti-HA (α-HA) antiserum or antibodies specific for the hypoxia markers, ADH and PGB1. Ponceau S staining was used to confirm equal loading. For all plots, different letters indicate significant differences between conditions (*P* < 0.05).

Given that the increased stabilisation of ERFVII transcription factors in *prt6-1 big-2* was associated with enhanced levels of key hypoxia response genes and proteins (Fig. 2, Fig. 3), we hypothesised that the double mutant might be more resistant to hypoxia than *prt6*. Chlorophyll retention is a marker of hypoxia tolerance, and we found that *prt6-1 big-2* and *prt6-5 big-2* had enhanced chlorophyll levels compared to the respective single mutants following hypoxia treatment (Fig S4A, C), with similar seedling survival rates (Fig. S4B, D). Ectopic expression of RAP2.3 in *prt6-5*, where *ADH* levels were very markedly elevated, dramatically enhanced both chlorophyll content and survival of seedlings following hypoxia (Fig. S4E, F), consistent with the role of RAP2.3 as a positive regulator of hypoxia responses.

VRN2 was first defined as a key regulator of vernalisation, but also contributes to hypoxia stress survival, with the *prt6-1 vrn2-5* mutant exhibiting lower tolerance than *prt6-1* [26]. However, *VRN2* was not required for hypoxia gene expression, indeed, expression of *PGB1* was increased when *prt6-1* and *prt6-1 big-2* were combined with the *vrn2-5* loss of function allele (Fig. 3E, F), suggesting that VRN2 may suppress expression of some hypoxia responsive genes under conditions where ERFVIIs are stabilised.

### The root transcriptome is extensively remodelled in *prt6-5 big-2* mutants

To obtain further insight into the impact of *BIG* on the PRT6/N-degron pathway and potentially other processes, we conducted mRNA sequencing (RNA-seq) analysis of *big-2, prt6-5, prt6-5 big-2* and Col-0 roots. One-centimetre root sections containing the root tip were selected to minimise potential developmental effects associated with the small size of *big-2* seedlings. As principal component analysis indicated that samples clustered tightly by genotype (Fig. S5A), we generated lists of differentially expressed genes (DEGs) for each mutant relative to wild-type, with a cut-off fold-change of 2 and adjusted p-value<0.01 (Dataset S2). Analysis of DEGs identified 92 and 119 genes up-regulated in *prt6-5* and *big-2*, respectively, with only 15 common DEGs. 341 and 438 genes were down-regulated in *prt6-5* and *big-2*, respectively, with 186 common to both data sets (Fig. 4A, B). Greater numbers of DEGs were identified in *prt6-5 big-2* and in many cases the fold changes of the common DEGs were significantly elevated in the double compared to the respective single mutants (Fig. 4C), indicative of a genetic interaction between *prt6-5* and *big-2*. There was significant overlap between genes up regulated in *prt6-5* and *prt6-5 big-2* roots with previously published microarray data from *prt6* and *ate1/2* seedlings [15, 31], but little overlap between *big-2* DEGs and the published data for *prt6* and *ate1/2* (Fig. S5B-D). Gene ontology (GO) term analysis [73] for ‘Biological Process’ revealed that hypoxia-related terms were highly enriched in *prt6-5* and *prt6-5 big-2* up regulated genes, whereas ‘Photosynthesis’, ‘Glycolate and dicarboxylate metabolism’ and ‘Carbon metabolism’ were enriched in *big-2* up regulated DEGs (Fig. S6). All the 49 ‘core’ genes known to be induced across cell types by hypoxia in wild-type plants [74] were present in the full transcriptome data set, with 21 up regulated in *prt6-5*. Comparison with transcriptome data from [75] revealed further hypoxia-responsive genes that are constitutively up regulated in *prt6-5* and *prt6-5 big-2* roots (Fig. 4D). In agreement with q-RT-PCR and immunoblotting data (Fig. 3), the fold-change in expression was markedly enhanced in *prt6-5 big-2* relative to the *prt6-5* single mutant (Fig. 4E) indicating a synergistic enhancement that is consistent with the increased stability of N-degron substrates such as the ERFVIIs. Six of the seven known N-degron pathway substrates were represented in the RNA-seq data set, but only the hypoxia-responsive genes *HRE1* and *HRE2* were classed as differentially expressed as defined by the 2-fold change cut-off (Fig. S5E).

**Fig. 4.**
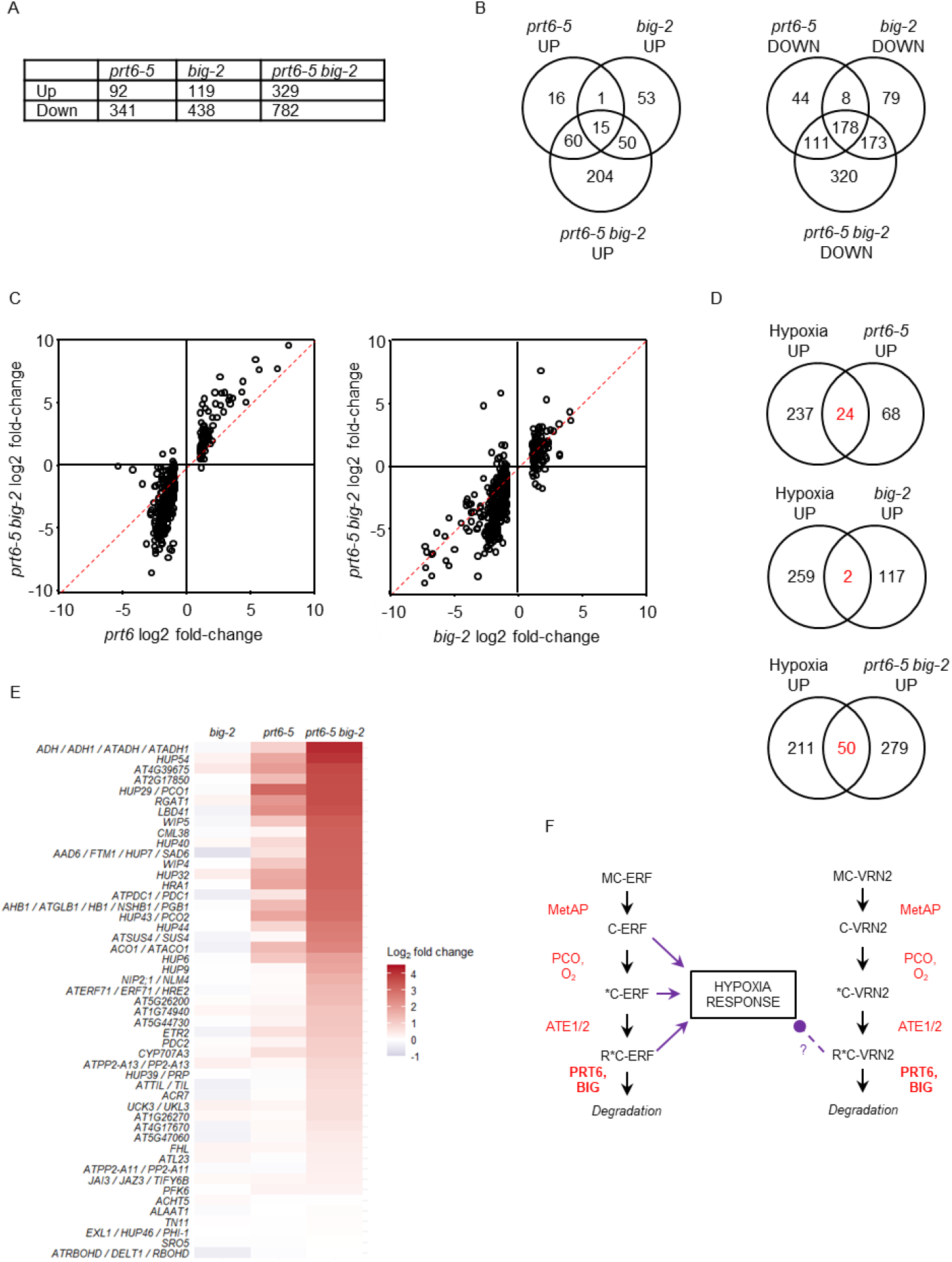
Impact of BIG on the root transcriptome. (A, B) Numbers of differentially expressed genes (DEGs) in roots of different mutant backgrounds. DEGs are defined as having a fold change greater than or equal to two at adjusted p-value<0.01. (C) Plots comparing the fold-change of transcripts in *big-2* and *prt6-5* single mutants with the double mutant, *prt6-5 big-2*. (D) Venn diagram showing overlap of DEGs with hypoxia responsive genes in Arabidopsis roots (differentially regulated following 7h of dark submergence [75]). (E) Heatmap showing fold change of 49 ‘core hypoxia’ genes [74] in the different mutant backgrounds. Gene names or AGI codes are shown to the left of the panel. (F) Scheme summarising impact of BIG on the N-degron pathway and the hypoxia response. Under normoxia, ERVII transcription factors and VRN2 are sequentially modified by methionine amino proteases (MetAP), plant cysteine oxidases (PCO) and arginyl-tRNA protein transferases (ATE1/2), such that the Nt Met is removed to reveal Cys2, which is oxidised (*C) and then arginylated (R*C). R*C-ERF and R*C-VRN2 are then targeted for degradation by PRT6 and also by a process involving BIG. Degradation is prevented in hypoxic conditions and in the absence of PRT6 (and BIG) function. The accumulation of ERFVIIs initiates the transcription of hypoxia responsive genes. The accumulation of VRN2 negatively influences the expression of certain hypoxia responsive genes. VRN2 also influences developmental processes as a component of the polycomb repressive complex 2 (not shown).

### Suberin deposition is repressed in *prt6* and *big-2* roots

Of the down-regulated genes, ‘Glucosinolate biosynthetic process’ was enriched in *prt6-5* and *prt6-5 big-2* DEGs, in agreement with previous findings for *ate1/2* [15] and ‘Cellular response to iron starvation’ was enriched in *prt6-5* and *big-2* (Fig. S7). Strikingly however, the most enriched terms for down-regulated genes in all genotypes were ‘Suberin biosynthetic process’ and ‘Cutin biosynthetic process’, two pathways which share common components [76] (Dataset S2). 39 genes associated with suberin biosynthesis and transport were down regulated in both *big-2* and *prt6-5*, accounting for almost all the steps in the pathway. These include long chain acyl-CoA synthetases, 3-ketoacyl CoA synthetases, fatty acid reductases, fatty acid omega-hydroxylases, glycerol acyltransferases, feruloyl acyltransferase, fatty alcohol caffeoyl-CoA transferase, 4-coumarate-CoA ligase, ABC transporters, lipid transfer proteins and GDSL lipases [76,77]. Fold changes of the differentially regulated genes were greater in *prt6-5 big-2*, compared to the respective single mutants (Fig. 5A). Moreover, six MYB transcription factors (*MYB9, MYB41, MYB53, MYB52, MYB93* and *MYB39/SUBERMAN*) which act in a hierarchical network to control suberin biosynthesis in Arabidopsis [78] are also down regulated in the N-degron pathway mutants, with fold-change increased in *prt6-5 big-2* relative to the single mutants (Fig. S8A). In agreement with this, 50 of 148 genes upregulated in *AtMYB41*-overexpressing plants [79] were down-regulated in *prt6-5 big-2* (Fig. S8B). Quantitative RT-PCR analysis showed that transcript levels of representative suberin genes were not significantly different between Col-0 and *prt6-1 rap2*.*12/2*.*2/2*.*3*, indicating that their repression in *prt6-1* roots requires RAP-type ERFVII transcription factors. In contrast, repression of suberin genes in *big-2* roots was ERFVII-independent (Fig. 5B). Finally, to explore the physiological significance of altered gene expression in the mutants, roots were stained with Fluorol Yellow 088. The suberised zone was less extensive in *prt6-5, big-2*, and *prt6-5 big-2* roots than in wild type Col-0 (Fig. 5C, D; Fig S8C). Taken together, the results indicate that suberin deposition is constrained in PRT6/N-degron pathway mutant roots by ERFVII stabilisation, and via an additional mechanism in *big-2*, pointing to shared and distinct roles for *BIG* and *PRT6* in control of this process.

**Fig. 5.**
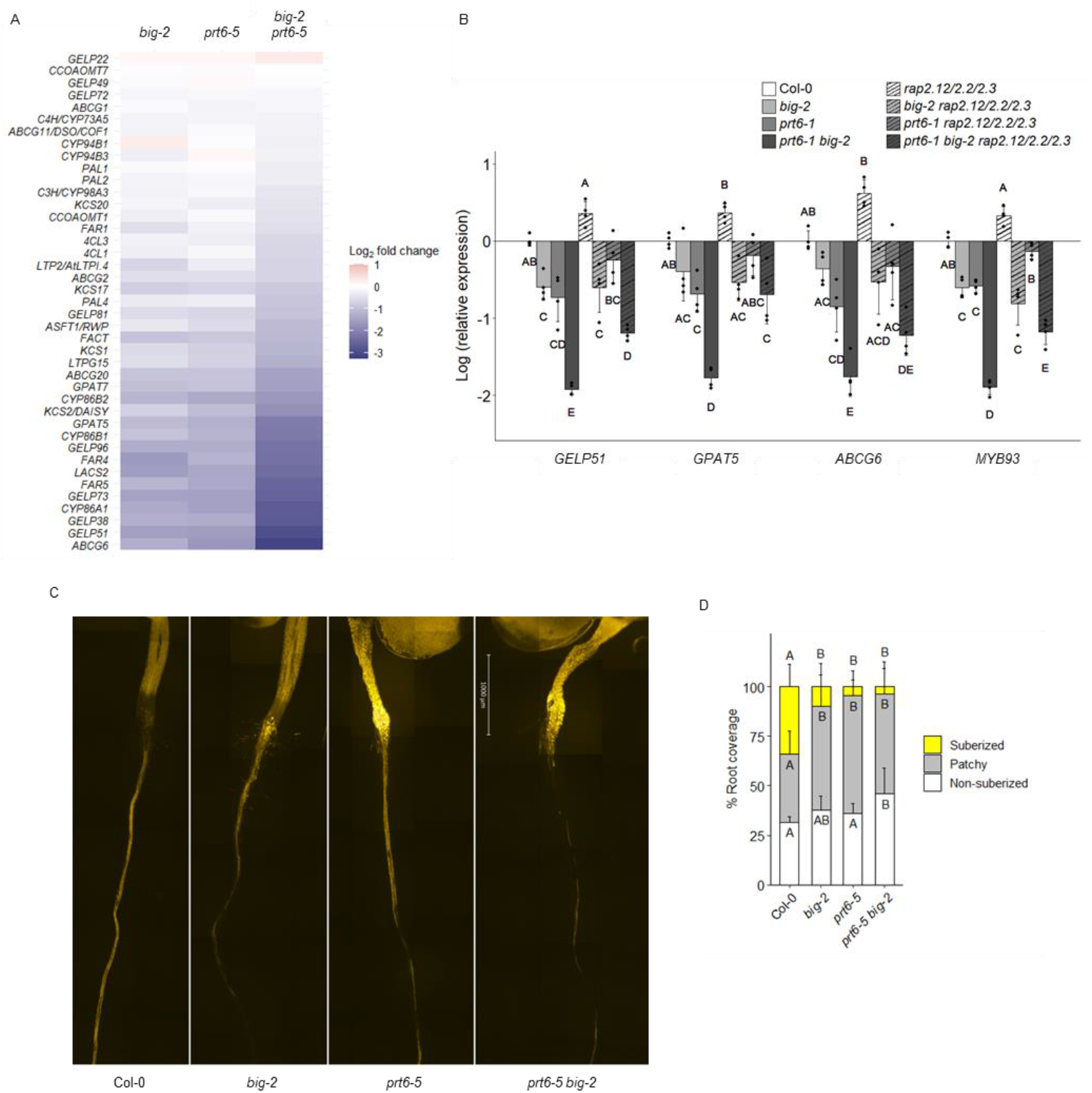
PRT6 and BIG regulate suberin deposition in roots. (A) Heatmap derived from RNA-seq data showing fold change of genes associated with suberin biosynthesis and deposition in the different mutant backgrounds. The gene list was curated from [76, 77, 81, 90]. Gene names or AGI codes are shown to the left of the panel. (B) Q-RT-PCR analysis of genes involved in suberin biosynthesis and deposition in 5-day old roots of wild type and mutants. Values are means ± SE (n=4); different letters indicate significant differences between genotypes (*P* < 0.05). (C) Representative composite micrographs showing Fluorol Yellow 088 staining of suberin in wild type and mutant roots (scale bar represents 1 mm). The full root image is shown in Fig. S8. (D) Quantification of suberin deposition along the root axis using three different zones: non-suberised, patchy, and continuous. Data are presented as percentage coverage of root length (n=10 roots; representative of two independent experiments); different letters indicate significant differences between genotypes for each region (*P* < 0.05).

## Discussion

Despite the physiological and agronomic importance of the plant Arg/N-degron pathways, not all of the molecular components have yet been identified [14]. Notably, incomplete stabilisation of the model substrate R-GUS in *Arabidopsis thaliana prt6* mutants [10] and the more severe phenotype of *ate1/2* compared to *prt6* [21] suggest the possibility of an additional N-recognin with specificity for PRT6/N-degrons. In this study, we provide several lines of evidence that the giant UBR box protein, BIG mediates turnover of proteins bearing type 1 N-degrons in concert with the PRT6/N-degron pathway and show that this influences the molecular response to low oxygen in Arabidopsis. Moreover, we demonstrate that BIG also contributes to the turnover of proteins with type 2 N-degrons via the PRT1/N-degron pathway.

Using N-degron pathway reporters, we demonstrated enhanced stability of a model PRT6 substrate, R-GUS as well as increased abundance of R-LUC and physiological substrates, HRE2 and VRN2 in the *prt6-5 big-2* mutant relative to *prt6-5* (Fig. 1; Fig. 2). The increased abundance of HRE2 and VRN2 was not driven by increased transcript, thus, whilst we cannot rule out increased translation, the data are consistent with increased stability in the double mutant background, as shown for R-GUS. It was not possible to test unequivocally whether RAP2.3 is similarly stabilised in *prt6-5 big-2* seedlings since double mutant plants expressing 35S::RAP2.3-HA did not set seed (Fig. S2). However, this observation, together with the partial rescue of delayed flowering and reduced fertility of *prt6-5 big-2* plants by genetic removal of *RAP* function (Fig. S3) strongly suggests that RAP2.3 is also hyperstabilised in the double mutant and that extreme stabilisation of RAP-type ERFVII transcription factors is deleterious to growth and reproduction.

Stabilisation of PRT6/N-degron pathway substrates in *prt6-5 big-2* plants markedly amplified the transcriptional response to hypoxia, as evidenced by q-RT-PCR, immunoblot, and especially RNA-seq analysis, and was accompanied by enhanced chlorophyll retention following hypoxia treatment (Fig. 2-4; Fig. S4). Higher order combination mutants demonstrated that BIG and PRT6 control the hypoxia response in seedlings exclusively through RAP-type ERFVII transcription factors (Fig 3 A-D). Interestingly, however, stabilisation of the PRC2 subunit VRN2 in PRT6/N-degron pathway mutants negatively influenced expression of the hypoxia-responsive gene *PGB1* (Fig. 4F). The mechanism by which this occurs remains to be explored but may involve methylation, given the known role of the PRC2 complex in epigenetic regulation. We did not explore the impact of enhanced VRN2 stabilisation on flowering since the *big-2* allele is in the Col-0 background which does not require vernalisation. Furthermore, ectopic expression of *VRN2* does not remove the requirement for vernalisation in ecotypes that require prolonged winter to initiate flowering [28].

RNA-seq analysis revealed that BIG and PRT6 not only play a role in the hypoxia response but also influence the expression of several other groups of genes, particularly a regulon associated with suberin biosynthesis, where *big-2* and *prt6-5* had an independent, partially additive negative effect on transcript abundance (Fig. 5; Fig. S8). In agreement with the lower expression of key *MYB* transcription factors and their downstream targets, suberin deposition was reduced in *big-2, prt6-5*, and *prt6-5 big-2* roots. Intriguingly, while *RAP2*.*12, 2*.*2* and *2*.*3* were required for the repression of suberin biosynthetic genes in *prt6*, repression in the *big-2* single mutant was independent of ERFVII transcription factors (Fig. S5B), suggesting that BIG influences other factors that control suberin deposition independently of PRT6. Suberisation is strongly influenced by hormones, including auxin which is associated with the growth phenotype of *big* alleles [48, 49, 50, 52, 54] and which has complex effects on suberin synthesis and degradation in the endodermis in Arabidopsis [79, 80]. It is tempting to speculate that dysregulation of auxin synthesis and transport underpin the reduced suberisation in *big-2*, however, there was no significant enrichment in transcripts related to auxin signalling pathways in *big-2* roots, in contrast to a previous transcriptome analysis employing leaves of a different *big* allele [63]. Whilst it is possible that there are tissue-specific differences in auxin-related gene expression that are not detected in the bulk root transcriptome, other mechanisms regulating suberin in *big-2* roots cannot yet be ruled out.

Taken together, our study provides evidence that BIG not only participates in the N-degron pathways, impacting different aspects of plant physiology, but likely also influences other processes. This raises interesting mechanistic questions regarding the operation of BIG in N-degron and possibly other proteostatic pathways. BIG contains numerous protein-protein interaction domains (Fig. S9; [48]) providing a platform for interaction with diverse protein partners and substrates. Proximity labelling identified both PRT6 and BIG as potential R-TurboID-interacting proteins (Fig. 1F), suggesting that BIG (like PRT6) may bind Arg/N-degrons, although a mutually compatible hypothesis is that BIG exists in a complex with PRT6 (see below). Reporter experiments revealed that BIG also works in concert with the PRT1 E3 ligase to mediate degradation of F-GUS (Fig. 1). Thus, BIG likely acts as an N-recognin for both type 1 and type 2 substrates. The mammalian N-recognins, UBR1 and 2 bind type 1 substrates via the UBR box and type 2 substrates at the Clp-S-like N-domain [82]. BIG, UBR4 and PRT6 each contain UBR boxes but lack the N-domain. Although the UBR box of PRT6 binds type 1 degrons, the N-degron binding sites of BIG and UBR4 have not been experimentally defined [7, 8, 10, 82, 83, 84]. PRT1 also lacks a ClpS-like domain and may recognise type 2 substrates via a ZZ domain, which is also present in BIG [12].

It remains to be determined whether BIG possesses intrinsic E3 ligase activity, since, like UBR4, it does not contain a canonical E3 ligase Really Interesting New Gene (RING) or HECT domain. Interestingly, regulatory proteasome subunits and the HECT E3 ligases, ubiquitin protein ligase (UPL)1, UPL2 and UPL3 were also enriched in R-TurboID samples, which may indicate the presence of an N-recognin/E3 ligase complex at the proteasome. In agreement with this, BIG co-purified with proteasome subunits and UPL1/3 in transiently transfected *Nicotiana benthamiana* [60] and UBR4 is present at the proteasome at significant, but sub-stoichiometric amounts in mammals [85, 86]. These observations are also consistent with previous reports of E3 ligases associated with the proteasome, including HECT E3 ligases [72] and yeast UBR1 [87]. In yeast, the HECT E3 ligase UFD4 binds UBR1 and increases processivity of polyubiquitination [88]; similarly in plants, substrates from diverse E3 ligases are relayed to UPLs which prevent substrate stalling at the proteasome [72]. Given that UBR4 interacts with a diverse array of protein partners including E2 ubiquitin conjugating enzymes and E3 ligases to degrade both N-degron pathway substrates and other protein targets [43, 45, 89], it is plausible that BIG serves as a versatile recognition component of the Arg/N-degron pathways and possibly other proteostatic mechanisms, interacting with one or more E3 ligases to mediate proteasomal degradation of a broad range of substrates (Fig. S10). Intriguingly, neither R-substrates nor F-substrates were stabilised in the single *big-2* mutant, suggesting that PRT6 and PRT1 are the dominant N-recognins *in planta*. It is possible that BIG may plays a more general role to prevent the release of potentially toxic, partly degraded proteins from the proteasome, recognising them via their neo-N-termini [85]. BIG may also participate in autophagic pathways in as is the case for UBR4 [9].

In conclusion, we have demonstrated that BIG participates in the Arg/N-degron pathways contributing to the turnover of ERVII transcription factors and VRN2 in the context of oxygen signalling, but that this does not underpin all of the known phenotypes associated with loss of *BIG* function. Key challenges for future work will be to identify novel substrates and E3 ligases associated with BIG and link them to its physiological functions.

## Materials and Methods

Genetic materials are listed in Table S1. Construction of combination mutants and reporter lines is described in *SI Appendix, SI Materials and Methods*. Where N-degron pathway reporters were compared in different genetic backgrounds, all lines were generated by crossing of a specific transgenic line, such that all genotypes within a given experiment contain the same transgene event.

## Acknowledgments

We thank Daniel Gibbs and Michael Holdsworth for sharing genetic materials, Kirsty Hassall for statistical advice, David Hughes for bioinformatic advice, and Graham Shephard for photography. Proteomics analyses were performed by the Mass Spectrometry Facility at Max Perutz Labs using the VBCF instrument pool. Work at Rothamsted Research was funded by the Biotechnology and Biological Sciences Research Council (BBSRC) through the Tailoring Plant Metabolism for the Bioeconomy Institute Strategic Grant BBS/E/C/000I0420 and the Green Engineering Institute Strategic Grant BB/X010988/1. Work in A.B.’s lab was supported by grant F7904B from the Austrian Research Agency FWF. Work in E.G.’s lab was supported by grant 13/IA/1870 from Science Foundation Ireland and by an Irish Research Council PhD scholarship (GOIPG/2017/2) to B.C.M.

## Notes

https://www.ebi.ac.uk/pride/

https://www.ncbi.nlm.nih.gov/

### Competing Interest Statement

The authors have declared no competing interest.

