## Supplementary material for "BIG participates in the Arg/N-degron pathways and the hypoxia response in *Arabidopsis thaliana*"

**This PDF file includes:**

Supplementary text
Figures S1 to S11
Tables S1 to S2
Legends for Datasets S1 to S2
SI References

**Other supplementary materials for this manuscript include the following:**

Datasets S1 to S2

**Supplementary Information Text**

### Extended Materials and Methods

#### Plant material

All genetic material used in this study is listed in Table S3. N-degron pathway mutants *prt6-5, prt6-1,* and *ate1/2* were crossed to *big-2 to* generate the double mutants *prt6-5 big-2*, *prt6-1 big-2*, and *big-2 ate1/2* triple mutant. N-degron pathway mutant alleles expressing DHFR-Ub-X-GUS reporter lines [1], X-LUC reporter lines [2,3], and pVRN2::VRN2:GUS [4] were crossed to *big-2* or *prt6-1 big-2* and 35S:HRE2-HA in Col-0 [5] was crossed to *prt6-5 big-2* and segregated into different backgrounds. 35S:RAP2.3-HA in Col-0 [6] was crossed to *big-2* and *prt6-5*, respectively, the resultant lines were crossed to each other, and seeds were maintained as 35S:RAP2.3-HA *prt6-5-/-big+/-.* Higher order loss of function mutants were obtained by crossing *rap2.12 rap2.2 rap2.3*, *prt6-1 rap2.12 rap2.2 rap2.3* [6], and *vrn2-5 prt6-1* [4], to *big-2* and *prt6-1 big-2*. All material was validated by PCR- or CAPS-based genotyping. Details of primers are given in Table S4.

Constructs for proximity labelling are based on vector R4 GWB601 [7], obtained from Addgene, and transformed into Col-0. The vector for R-Turbo (2548_HpaI_Turbo-NESYFP) is shown in Fig. S11, its M-Turbo counterpart differs by only two bases (exchange ATG for AGA, codon for first amino acid after ubiquitin cleavage).

R-LUC reporter lines in a wild-type Col-0 background [3]; based on constructs generated by [2] were crossed with *prt6-5*, *big-2* and *prt6-5 big-2* mutants. Lines containing the R-LUC reporter were selected on 0.5x MS + 0.5% (w/v) sucrose, 0.8 g/L agar plates containing 20 mg/L Basta, and subsequently (i) genotyped to isolate homozygous mutants for *big-2* and *prt6-5*; and (ii) sequenced to confirm the identity of the R-LUC reporter.

#### Growth of Arabidopsis

Seeds were raised from plants grown in Levington's F2S compost under long day conditions (16 h day/8 h night; 23°C/18°C); all genotypes to be compared were raised in the same controlled environment cabinet. Seeds were harvested, sieved (< 425 µm; Endecotts, London, UK) and stored at room temperature. After-ripened seeds were surface-sterilised and sown on 0.5X Murashige and Skoog (MS) medium containing 0.5% (w/v) sucrose and 0.8% (w/v) plant agar (Duchefa). After 2-3 d dark chilling at 4°C, plates were grown in long days (16 h/8 h; 22°C) for 4 to 6 d.

#### Genotyping

For DNA isolation, frozen tissue was homogenized using a GenoGrinder (1750 rpm for 1.5 min) equipped with metal blocks pre-chilled with liquid $N_2$. 500 µL pre-warmed CTAB buffer [2% cetyl trimethylammonium bromide, 1% polyvinyl pyrrolidone (MW = 40,000), 1.4 M NaCl, 0.1 M Tris HCl, 20 mM EDTA, pH 5.0] was added to the powder, and incubated at 60°C for 30 min. Samples were centrifuged at 10,000 x g for 5 min, 5 µL RNase-A (10 mg/mL) was added to the supernatant and incubated for 15 min at room temperature. DNA was extracted by addition of an equal volume of chloroform/isoamyl alcohol (24:1 v/v). Following centrifugation at 13,000 x g for 1 min, DNA in the upper aqueous phase was precipitated by adding 0.7 vol of isopropanol and incubating at -20°C for 15 min. DNA was pelleted by centrifuged at 13,000 x g for 10 min and the pellet washed twice with 400 µL pre-chilled 70% ethanol and dried briefly, before dissolving in 100 µL TE buffer (10 mM Tris, 1 mM EDTA, pH 8.0). PCR was performed using a 20 µL total reaction volume, consisting of: 1X DreamTaq Green PCR Master Mix, 500 nM Forward Primer, 500 nM Reverse Primer, 10% v/v plant genomic DNA. Primers used are given in Table S4.

### GUS staining

Seedlings grown on sterile agar medium were immersed in 1 mL GUS assay buffer [100 mM sodium phosphate buffer (pH 7.0), 0.1 % Triton X-100, 0.5 mg/mL X-GlucA, 500 µM potassium ferricyanide, 500 µM potassium ferrocyanide] in sterile 24-well plates, vacuum infiltrated for 30 min in darkness, then wrapped in foil and incubated at 37 °C overnight. Chlorophyll was removed by incubation in 85% ethanol, 15% acetic acid with gentle agitation for 2-4 h until cleared, after which the seedlings were placed in sterile water. Seedlings were arranged onto agar plates for photography.

### X-LUC assays

Seedlings stably expressing the R-LUC N-degron reporter construct were grown vertically on 0.5x MS + 0.5% (w/v) sucrose and 0.8 g/L agar plates containing 20 mg/L Basta to select for the presence of the reporter. Plates were kept at 4°C in the dark for 3 days and then transferred to continuous light at 19.5°C for 7 days. Forty seedlings per genotype were harvested and immediately frozen in liquid nitrogen. Frozen tissue was ground using a drill and pestle and the powder was split equally between two tubes for (i) LUC enzymatic assays and (ii) RNA extraction followed by q-RT-PCR to normalise the LUC enzymatic activities to the expression of the *LUC* gene in each of the samples.

To test the enzymatic R-LUC activity, proteins were extracted from frozen ground tissue using 1x Luciferase Cell Culture Lysis Reagent (CCLR) (Promega), supplemented with 1 mM phenylmethylsulfonyl fluoride (PMSF) and 1:100 plant Protease Inhibitor Cocktail (Sigma-Aldrich). Samples were centrifuged at 12,000 x g for 10 minutes at 4°C to pellet cellular debris. Protein concentration was determined using the Bradford protein assay. Enzymatic LUC activity was measured as described in [3, 8]. Briefly, CCLR protein extract (1 µL) was added to 100 µL LAR buffer (20 mM tricine, pH 7.8, 1.07 mM $(MgCO_3)_4.Mg(OH)_2.5H_2O$, 2.67 mM $MgSO_4$, 0.1 mM ethylenediaminetetraacetic acid (EDTA), 33.3 mM dithiothreitol (DTT), 270 µM coenzyme A, 470 µM luciferin, 530 µM ATP) in a 96-well plate (Sterilin). Luminescence was measured using a POLARstar Omega microplate reader (BMG LABTECH) for 10 seconds.

To determine expression levels of the R-LUC reporter, total RNA was extracted using the Spectrum Plant Total RNA Kit (Sigma Aldrich/Merck) according to the manufacturer's instructions. Reverse transcription reactions were set up using 1000 ng of total RNA, RevertAid Reverse Transcriptase (Thermo Fisher) and associated buffer, RiboLock RNase inhibitor (Thermo Fisher), oligo(dT)18 and 1 mM dNTP mixture at 42°C for 45 minutes. qPCR reaction mixtures were prepared in LightCycler 480 96-well plates (Roche) with 1 µL of cDNA, 1 µL of primer pair mixture (1 µM final concentration each primer; Table S4), 5 µL 2x SYBR green master mix (Roche), with nuclease-free water added to a final volume of 10 µL per well. qPCR reactions were carried out in a LightCycler 480 instrument (Roche). The second derivative maximum method was used to determine crossing point (Cp) values.

### Immunoblotting

Six-day-old roots or seedlings were harvested. Protein extraction and immunoblotting were performed as described in [9], with the exception that 1% BSA in PBS-T was used as the blocking agent in the case of the anti-Biotin blots. Briefly, proteins were separated in precast 4–12% Bis-Tris gels using 1X SDS MES buffer and transferred to polyvinylidene fluoride using iBlot™ 2 Dry Blotting System (ThermoFisher, Waltham, MA, USA). Primary antibodies were used at the following dilutions: ADH (Agrisera, Sweden), 1:3000; PGB1 (Hartman 2018) 1:3000, GUS (Sigma-Aldrich; G5420), 1:1000; HA (H 3663, Sigma) 1:1000 and biotin (Sigma, BN-34) 1:2,000. The secondary antibodies used were anti-rabbit horseradish peroxidase conjugate (A0545; Sigma) diluted 1:50,000 (for ADH, PGB1 and GUS), m-IgGk BP-HRP (sc-516102; Santa Cruz Biotechnology) diluted 1:15,000 (for HA) or anti-mouse (GE Healthcare NA931) diluted 1:10,000 (for biotin). Blots were then washed and developed with SuperSignal™ West Pico PLUS Chemiluminescent Substrate (Thermo Fisher Scientific).

**Real time quantitative reverse-transcription PCR (RT-qPCR)**

Six-day-old roots or seedlings were harvested, and total RNA was extracted using an RNeasy Plant Mini Kit (Qiagen) and treated using a TURBO DNA-free™ Kit (Invitrogen™), or a Monarch® Total RNA Miniprep Kit (New England Biolabs, Inc), with on-column DNAse I treatment. A RevertAid™ First Strand cDNA Synthesis Kit (Thermo Scientific) and anchored -oligo(dT)18 were used for cDNA synthesis for a two-step RT-PCR. SYBR® Green JumpStart™ Taq ReadyMix™ was used for real-time PCR using a Lightcycler® 96 Instrument (Roche) or a Quantstudio 6 Pro (Thermo), according to manufacturers' instructions. Three to four biological replicates were included for each genotype, with two technical replicates per sample. Relative quantification was performed using both *ACT2* (At3g18780.2) and *TUB4* (At5g44340.1) as references. For the experiments presented in Fig. 3C and 3E, a single reference gene was used (*ACT2*) due to practical constraints. *UBQ10* and AT5G18800 were used as references for the experiment presented in Fig. S8D. Relative gene expression was calculated using the $2^{-\Delta\Delta Ct}$ method  Primers used are given in Table S4.

**RNA-seq**

After-ripened seeds were surface-sterilised and plated on nylon mesh (Sefar NITEX, 03-110/47; Heiden, Switzerland) overlaid on 0.5X Murashige and Skoog (MS) medium containing 0.5 % sucrose and 0.8% plant agar. One cm sections containing the root tip were harvested and RNA was extracted using an RNeasy Plant Mini Kit (Qiagen) and treated using a TURBO DNA-free™ Kit (Invitrogen™). RNA sequencing and data analysis was done using Illumina HiSeq (2 x 150 paired end reads) by Genewiz. Briefly, sequence reads were trimmed to remove possible adapter sequences and nucleotides with poor quality using Trimmomatic v.0.36. The trimmed reads were mapped to the *Arabidopsis thaliana* TAIR10 reference genome available on ENSEMBL using the STAR aligner v.2.5.2b. Unique gene hit counts were calculated by using feature Counts from the Subread package v.1.5.2. Only unique reads that fell within exon regions were counted. Since a strand-specific library preparation was performed, the reads were strand-specifically counted. Differential expression analysis was performed using DESeq2. The Wald test was used to generate p-values and log2 fold changes. Genes with an adjusted p-value < 0.05 and absolute log2 fold change > 1 were called as differentially expressed genes for each comparison. The RNA-seq data files for this study have been uploaded to NCBI (https://www.ncbi.nlm.nih.gov/) under project number PRJNA975350, with accession numbers SAMN35345055-SAMN35345074.

**Hypoxia assays**

Hypoxia conditions were generated by anaerobic atmosphere generation bags (68061 Sigma) in an anaerobic jar (28029 Sigma) according to the manufacturer's instructions. Seedlings were treated with hypoxia in the dark by enclosing the plates in an anaerobic jar, from which oxygen was reduced to below 1% within 1 h, monitored by smart sensor Oxygen Detector AR8100. Controls were kept in the dark for the same period of time under normal oxygen conditions. After 3 d recovery in the light, seedlings were photographed, weighed, submerged in 80% acetone overnight at 4°C, shielded from light, to extract chlorophyll. Absorbance at 646 nm and 663 nm was used to estimate total chlorophyll [10]. For survival scoring, seedlings were visually assessed and assigned a score based on their appearance as in [5]: 1 for no remaining chlorophyll, 3 for partial chlorophyll coverage, and 5 for complete chlorophyll coverage. Scores were aggregated to produce a mean survival score for each plate of seedlings.

**Statistical analysis**

Statistical analyses were performed using the R environment. For analysis of multiple genotypes and/or transgenic lines, one-way ANOVA followed by Tukey's multiple comparison test was used, after log transformation where indicated by linear model fitting.

**Proximity labelling**

Arabidopsis plants (15 seedlings per well, 24 well plate) were grown in liquid culture (1ml 1x MS medium with 1% sucrose per well) for 1 week under long day conditions. Medium was exchanged with medium supplemented with 10 µM Bortezomib and 50 µM biotin 75 min before harvest. Plants were washed 4x with 2 ml ice cold water, then 1 ml ice cold water was added before they were dried and snap frozen in liquid $N_2$. Tissue was homogenised in a pre-cooled Tissue Lyser (2x 10 min, 28 Hz) and 170µl extraction buffer [50 mM Tris pH 7.5; 150 mM NaCl; 1 mM EDTA; 0,5% NP-40 substitute; 3 mM DTT; plant protease inhibitor cocktail (Sigma)] added. Extracts from 3 wells were pooled (= ~500 µl crude extract as input). After centrifugation at 4°C, 500 µl of the supernatant was loaded onto a Sephadex G-25 column (Cytiva MiniTrap PD-10) and eluted with 1 ml extraction buffer to remove free biotin. The protein concentration of the eluate was determined by Bradford assay and amounts of extracts were adjusted to the sample with the lowest protein concentration (2.3 mg total protein). Samples were incubated with Pierce Magnetic Streptavidin beads equilibrated in extraction buffer (25µl beads per sample) for 60 min with rotation at 4°C. The beads were washed 4x with 1ml wash buffer (20 mM Tris pH 7.5; 500 mM NaCl; 0,5 mM EDTA), transferred to a fresh low binding tube, washed again with 1ml wash buffer and finally resuspended in 500µl wash buffer. Three technical replicates per genotype were submitted to proteomic analysis. Two additional biological replicates (each including technical replicates) gave similar results.

**Sample preparation for mass spectrometry analysis**

The beads were resuspended in 50 µL 1 M urea and 50 mM ammonium bicarbonate. Disulfide bonds were reduced with 2 µL of 250 mM dithiothreitol (DTT) for 30 min at room temperature before adding 2 µL of 500 mM iodoacetamide and incubating for 30 min at room temperature in the dark. The remaining iodoacetamide was quenched with 1 µL of 250 mM DTT for 10 min. Proteins were digested with 150 ng LysC (mass spectrometry grade, FUJIFILM Wako chemicals) in 1.5 µL 50 mM ammonium bicarbonate at 25°C overnight. The supernatant without beads was digested with 150 ng trypsin (Trypsin Gold, Promega) in 1.5 µL 50 mM ammonium bicarbonate followed by incubation at 37°C for 5 hours. The digest was stopped by the addition of trifluoroacetic acid (TFA) to a final concentration of 0.5 %, and the peptides were desalted using C18 Stagetips [11].

**Liquid chromatography-mass spectrometry analysis**

Peptides were separated on an Ultimate 3000 RSLC nano-flow chromatography system (Thermo Fisher), using a pre-column for sample loading (Acclaim PepMap C18, 2 cm × 0.1 mm, 5 µm, Thermo Fisher), and a C18 analytical column (Acclaim PepMap C18, 50 cm × 0.75 mm, 2 µm, Thermo Fisher), applying a segmented linear gradient from 2% to 35% and finally 80% solvent B (80 % acetonitrile, 0.1 % formic acid; solvent A 0.1 % formic acid) at a flow rate of 230 nL/min over 120 min.

Eluting peptides were analysed on an Exploris 480 Orbitrap mass spectrometer (Thermo Fisher) coupled to the column with a FAIMS pro ion-source (Thermo Fisher) using coated emitter tips (PepSep, MSWil) with the following settings: the mass spectrometer was operated in DDA mode with two FAIMS compensation voltages (CV) set to -45 or -60 and 1.5 s cycle time per CV. The survey scans were obtained in a mass range of 350-1500 m/z, at a resolution of 60k at 200 m/z, and a normalized AGC target at 100%. The most intense ions were selected with an isolation width of 1.2 m/z, fragmented in the HCD cell at 28% collision energy, and the spectra recorded for max. 100 ms at a normalized AGC target of 100% and a resolution of 15k. Peptides with a charge of +2 to +6 were included for fragmentation, the peptide match feature was set to preferred, the exclude isotope feature was enabled, and selected precursors were dynamically excluded from repeated sampling for 45 s.

#### Proteomics data analysis

MS raw data split for each CV using FreeStyle 1.7 (Thermo Fisher), were analysed using the MaxQuant software package (version 2.1.0.0; [12]) with the Uniprot *Arabidopsis thaliana* reference proteome (version 2022.01 www.uniprot.org), target sequences, as well as a database of the most common contaminants. The search was performed with full trypsin specificity and a maximum of two missed cleavages at a protein and peptide spectrum match false discovery rate of 1%. Carbamidomethylation of cysteine residues was set as fixed, oxidation of methionine, and N-terminal acetylation as variable modifications. For label-free quantification the "match between runs" only within the sample batch and the LFQ function were activated - all other parameters were left at default. MaxQuant output tables were further processed in R 4.2.1 (https://www.R-project.org) using Cassiopeia_LFQ (https://github.com/moritzmadern/Cassiopeia_LFQ). Reverse database identifications, contaminant proteins, protein groups identified only by a modified peptide, protein groups with less than two quantitative values in one experimental group, and protein groups with less than 2 razor peptides were removed for further analysis. Missing values were replaced by randomly drawing data points from a normal distribution model on the whole dataset (data mean shifted by -1.8 standard deviations, a width of the distribution of 0.3 standard deviations). Differences between groups were statistically evaluated using the LIMMA 3.52.1 [13] at 5% FDR (Benjamini-Hochberg).  The mass spectrometry proteomics data have been deposited to the ProteomeXchange Consortium via the PRIDE [14] partner repository with the dataset identifier PXD041610.

#### Suberin quantification

Seeds were surface-sterilised and plated on 0.5x MS + 0.5% sucrose containing 0.8 % agarose. After 2–3 d dark chilling at 4°C, seedlings were grown vertically in long days (16 h/8 h; 22°C ) for 5 days and stained with Fluorol Yellow 088, as described in [15]. Tiled images were captured across whole seedlings using a Zeiss Axio Imager.Z2 microscope (10X objective and GFP fluorescence filters: excitation 450 – 490 nm; emission 500 – 550 nm, illumination 450 – 488 nm) and Zen3.0 blue edition software. Seedlings were initially viewed using brightfield imaging at a low light level to define the region to be scanned and create focal points ("support points") along the length of the root. Entire seedlings were then scanned using fluorescence contrast imaging with 100% light intensity (excitation 488 nm; emission 509 nm) and a 10% overlap of tile images for alignment and stitching. Images were pseudo-coloured using the "YellowToWhite" LUT, annotated with a 1000 µm scale bar, and exported as TIFF files at 70% of the original size. Suberisation patterns were quantified using ImageJ to measure the length of the different regions in µm: "suberised" for continuous suberisation, "patchy" for partial suberisation, and "non-suberised" for the region with no suberised cells. Results were expressed as the percentage of the total root length.

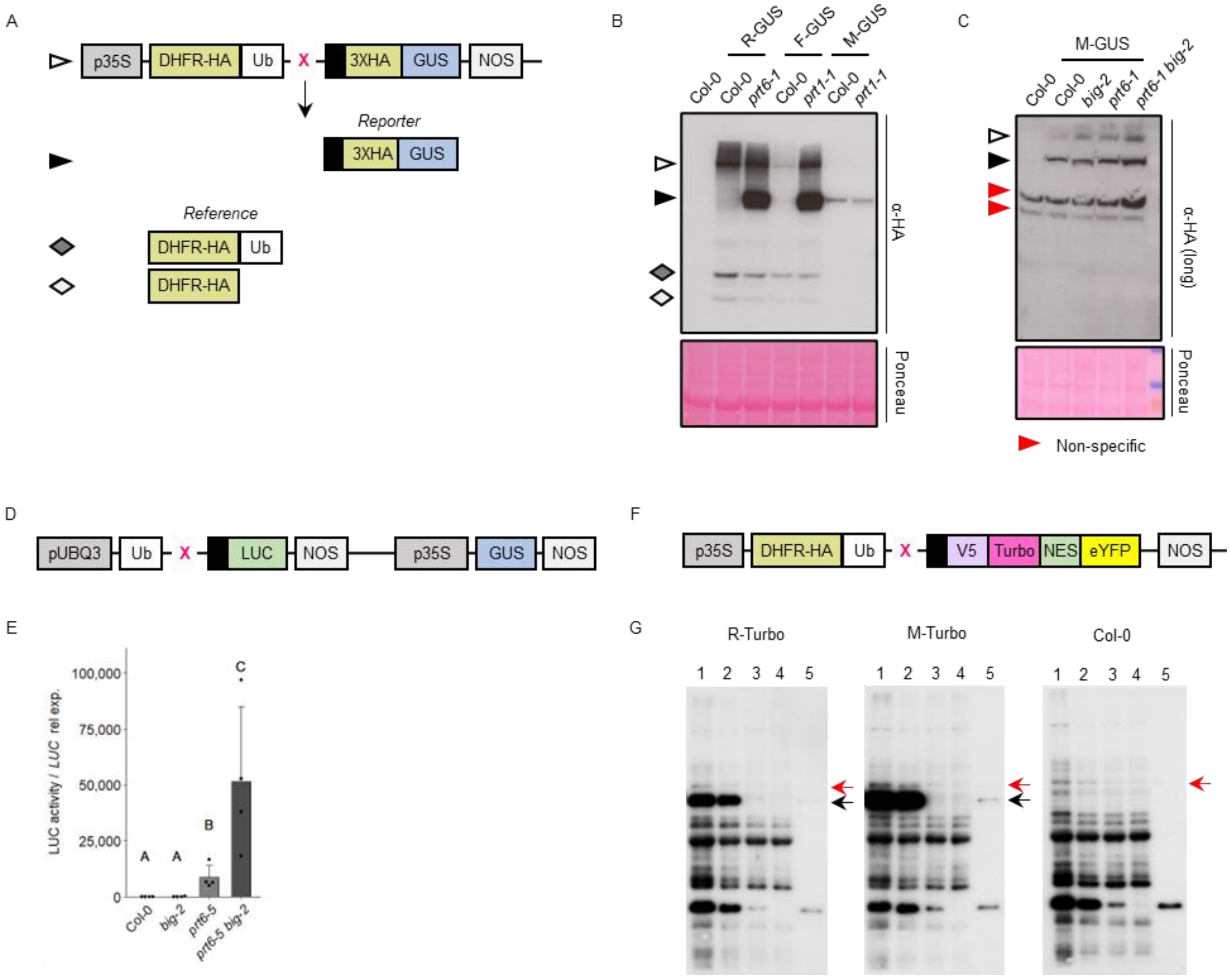

**Fig. S1. BIG influences the stability of model type 1 and type 2 Arg/N-degron pathway substrates.**

(A) Generation of N-degron pathway substrates. Constructs driven by the constitutive CaMV35S promoter (p35S) encode a fusion of mouse dihydrofolate reductase (DHFR, yellow) to ubiquitin (variant K48R; Ub, white), followed by Escherichia coli beta glucuronidase (GUS, blue). Ubiquitin-specific proteases remove ubiquitin co-translationally to release the GUS reporter protein and reveal a new N terminus (a variable residue, indicated by X). The GUS ORF is extended by unstructured amino acids (black) to enhance the effect of destabilising amino-terminal residues. The cleavage also creates a stable DHFR reference protein and HA epitopes enable immunological detection of both products [1]. (B) Detection of N-degron pathway substrates by immunoblotting crude protein extracts from 6-d old seedlings of different genotypes expressing X-GUS reporters. Symbols to the left indicate the protein products shown in (A). Ponceau S staining was used to confirm equal loading. (C)  As (B), for M-GUS. The blot was developed with a long exposure in order to detect the reporter protein. Expression of the M-GUS transgene is low in these lines, due to silencing, and it was not possible to detect the stable reference. Symbols to the left indicate the protein products shown in (A); other bands indicated by red symbols correspond to cross-reacting proteins also present in plants carrying no transgene. (D) Alternative system for generation of N-degron pathway substrates [2]. A fusion of Ub (white) to LUCIFERASE (LUC, green) is driven by the UBQ3 promoter (pUBQ3, grey). A short linker (represented by a black rectangle) is present between the variant residue X and the LUC reporter. In the same construct, GUS (blue) driven by the CaMV35S promoter (p35S) acts as a stable reference protein. NOS: nopaline synthase transcriptional terminator sequence. (E) Luciferase activity in 7-d old seedlings of different genotypes expressing R-LUC. Luciferase activity was normalised according to LUC transcript levels because the GUS normalising cassette was variably silenced during crossing, probably due to the presence of multiple copies of the CaMV35S promoter in the mutant backgrounds. Values are ± SD (n=4); different letters indicate significant differences between conditions ($P < 0.05$). (F) Construct used for proximity ligation assay. Turbo-NES-eYFP [7] was modified to harbour an N-terminal Ub fusion construct (DHFR-HA-Ub) followed by the N-terminal amino acid of choice in front of a linker sequence (black), expressed from the Ubiquitin 10 (pUBQ10) promoter. (G) Anti-biotin immunoblots to follow the enrichment of biotinylated protein in extracts from Col-0 and plants expressing R-Turbo and M-Turbo. Lane 1, crude extract; lane 2, extract after desalting; lanes 3 and 4, supernatant after two subsequent incubations with streptavidin beads (unbound fractions); lane 5, streptavidin beads after first incubation. Black arrows indicate the bait proteins R-Turbo and M-Turbo (70.2 kDa), and the red arrow indicates a non-specific background band.

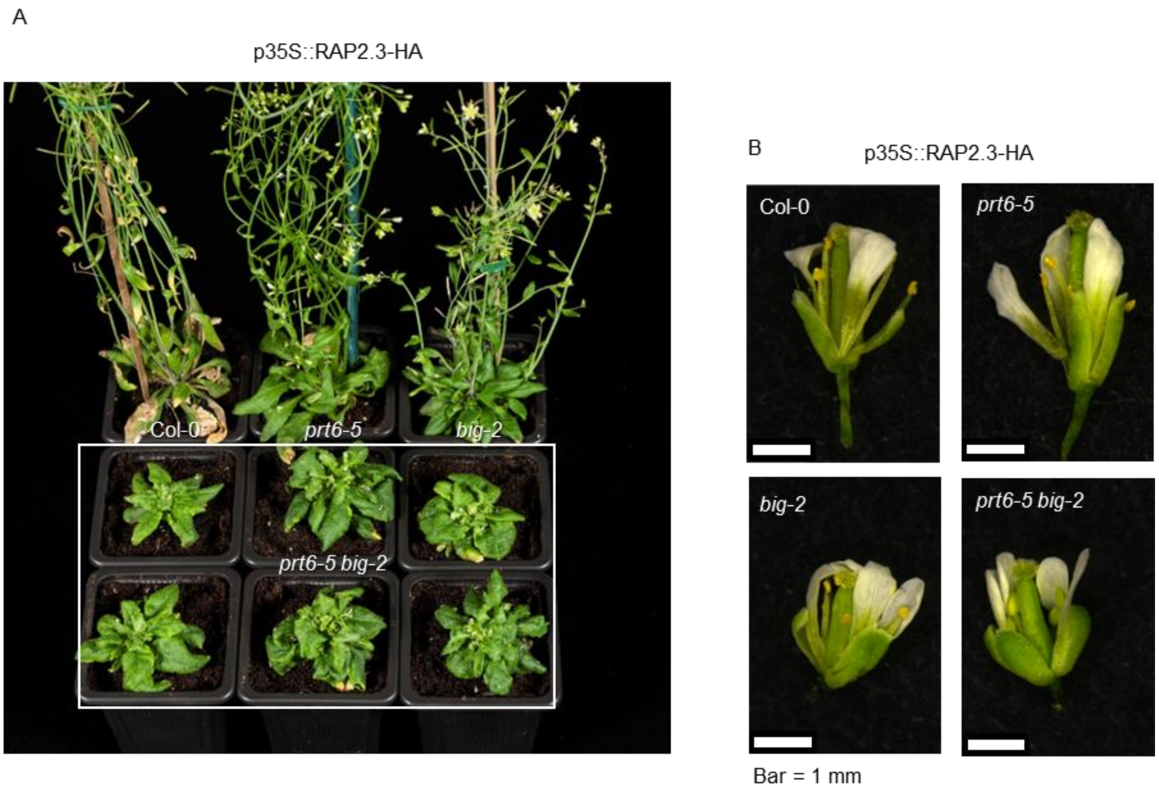

**Fig. S2. Morphology of plants ectopically expressing RAP2.3-HA.**

(A) Plants were germinated on 0.5 X MS containing 0.5 % sucrose, transplanted to soil, and photographed after 5 weeks under long day conditions. (B) Flowers from the primary bolt of different genotypes expressing p35S::RAP2.3-HA were partially dissected to reveal stamens and gynoecia. Bar = 1 mm.

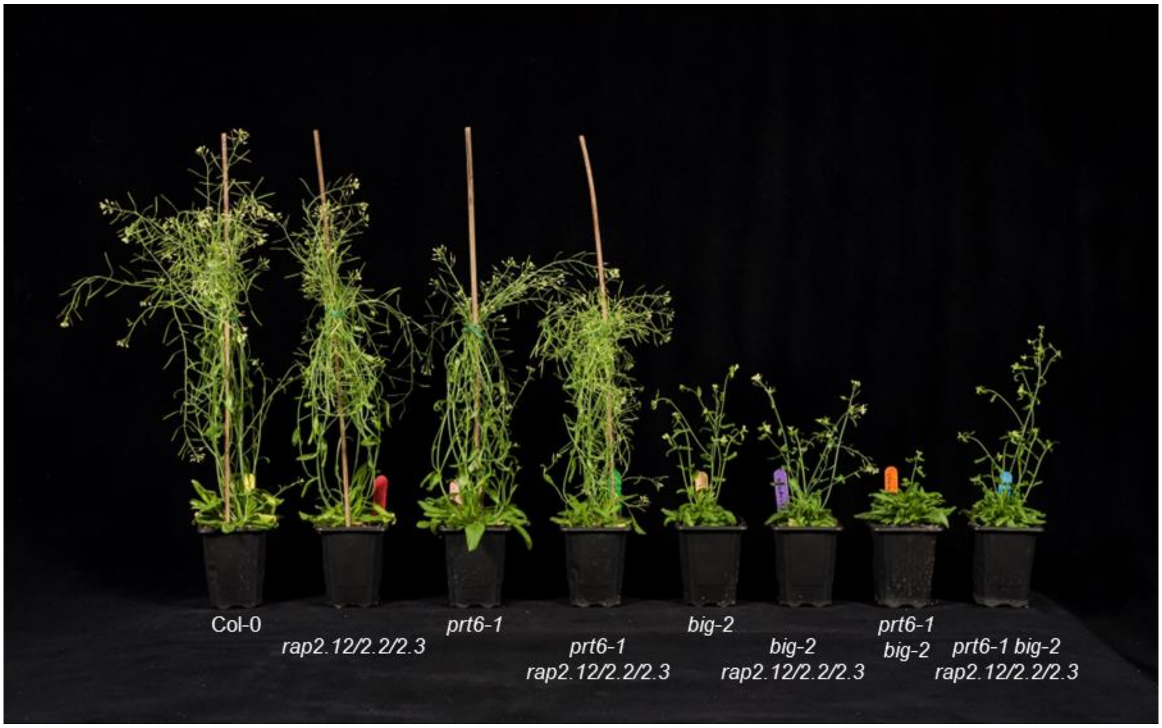

**Fig. S3 Morphology of PRT6/N-degron pathway and *erfvii* combination mutants.**
Plants were grown in soil for six weeks under long day conditions, prior to photography.

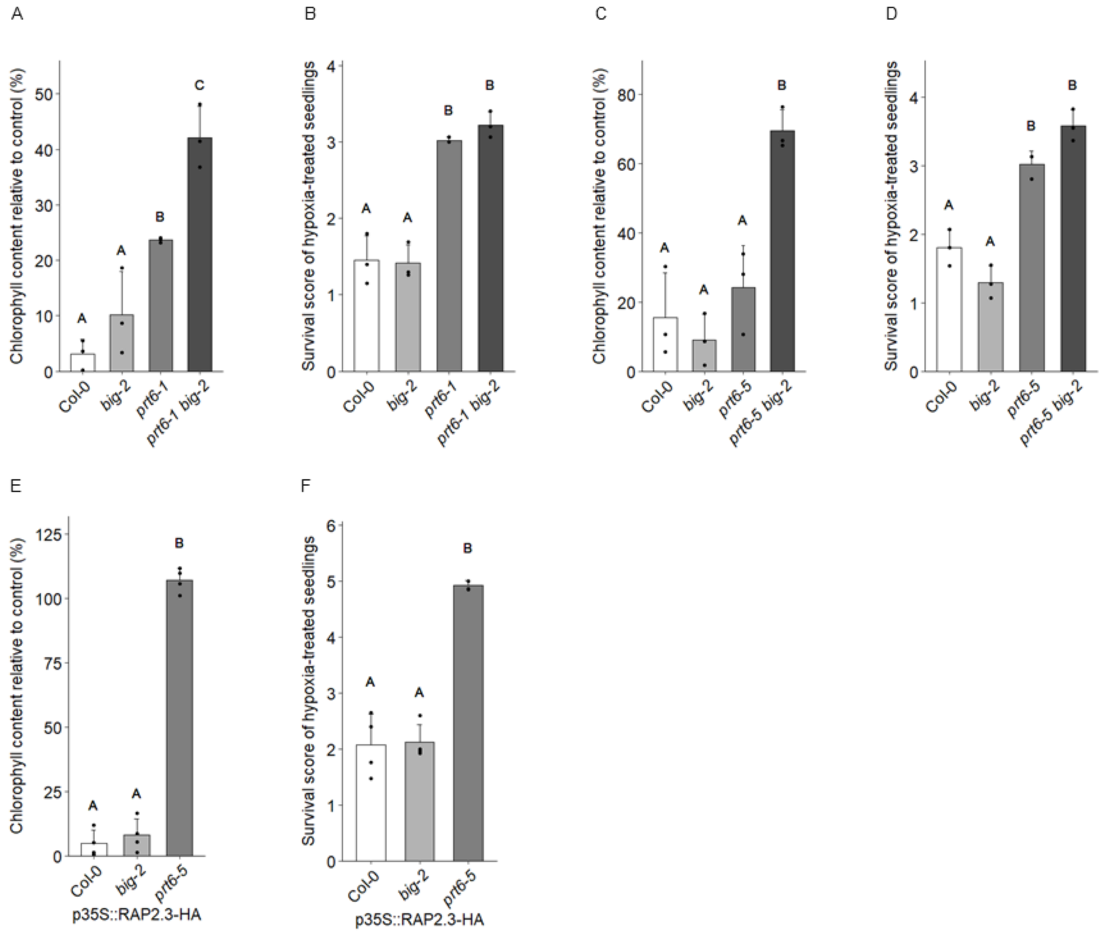

**Fig. S4 Hypoxia response of PRT6/N-degron pathway mutants.**

Seedlings of untransformed mutants (A-D) and transgenic lines ectopically expressing p35S::RAP2.3-HA (E,F) were grown on 0.5 X MS medium containing 0.5 % sucrose for 4 days, subjected to 5 h hypoxia, and analysed after 3 d recovery in the light. (A,C,E) Chlorophyll content of hypoxia treated seedlings relative to control; (B,D,F) Survival data. Values are means ± SD (n=3). For all plots, different letters indicate significant differences between conditions ($P < 0.05$).

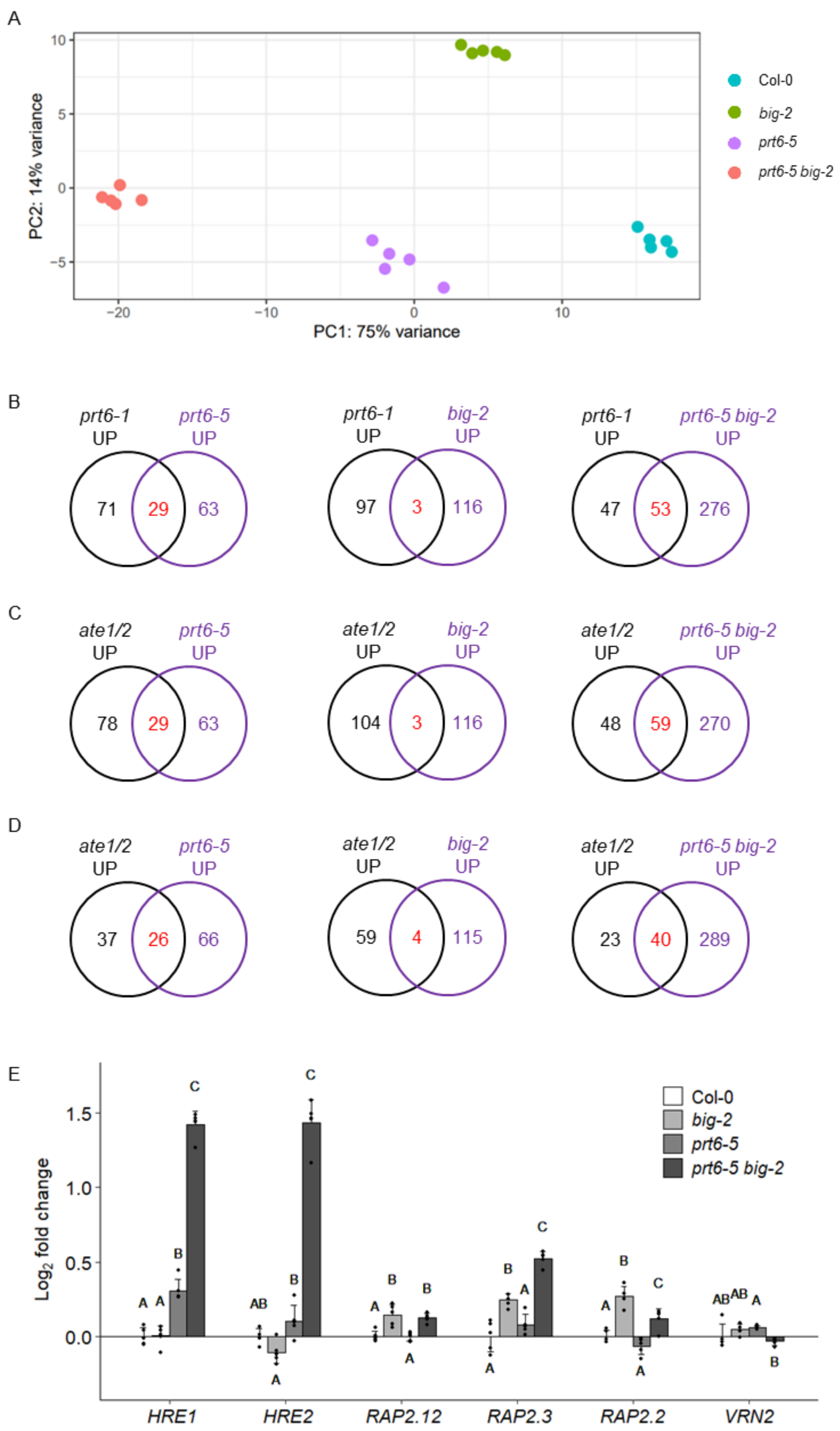

**Fig. S5 Transcriptome analysis of N-degron pathway mutants.**

(A) Principal component analysis of RNA-seq data. (B-D) Venn diagrams showing comparison of differentially expressed genes from the RNA-seq data reported in this study (purple) with differentially regulated genes in published microarray studies (black): (B) *prt6-1* seedlings [5], (C) *ate1 ate2* seedlings [5], (D) *ate1 ate2* seedlings [16]. (E) Expression of genes encoding known PRT6/N-degron pathway substrates in roots of different mutant backgrounds, relative to Col-0; different letters indicate significant differences between conditions ($P < 0.05$). *LITTLE ZIPPER2* (*ZPR2*) was not identified in the data set.

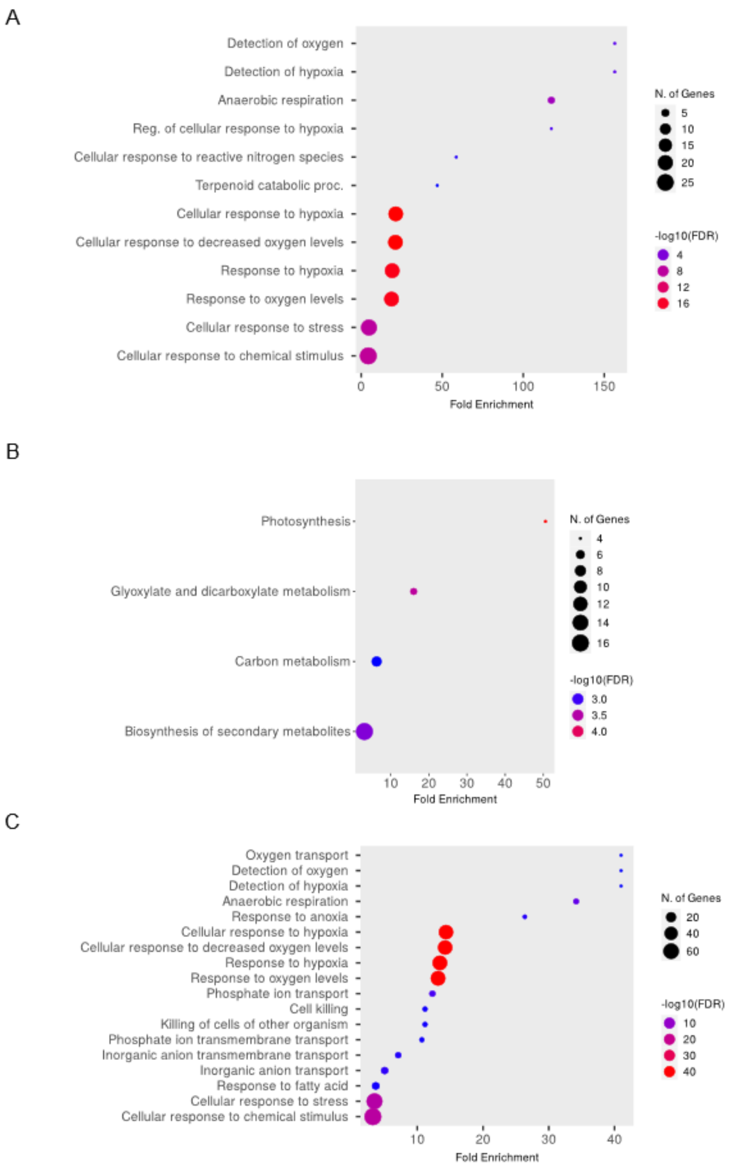

**Fig. S6 Gene Ontology terms enrichment in genes up-regulated in roots of N-degron pathway mutants.**

Differentially expressed gene lists were analysed with ShinyGO [17] using all genes identified in the RNA-seq data set as background. (A) *prt6-5*. (B) *big-2*. (C) *prt6-5 big-2*.

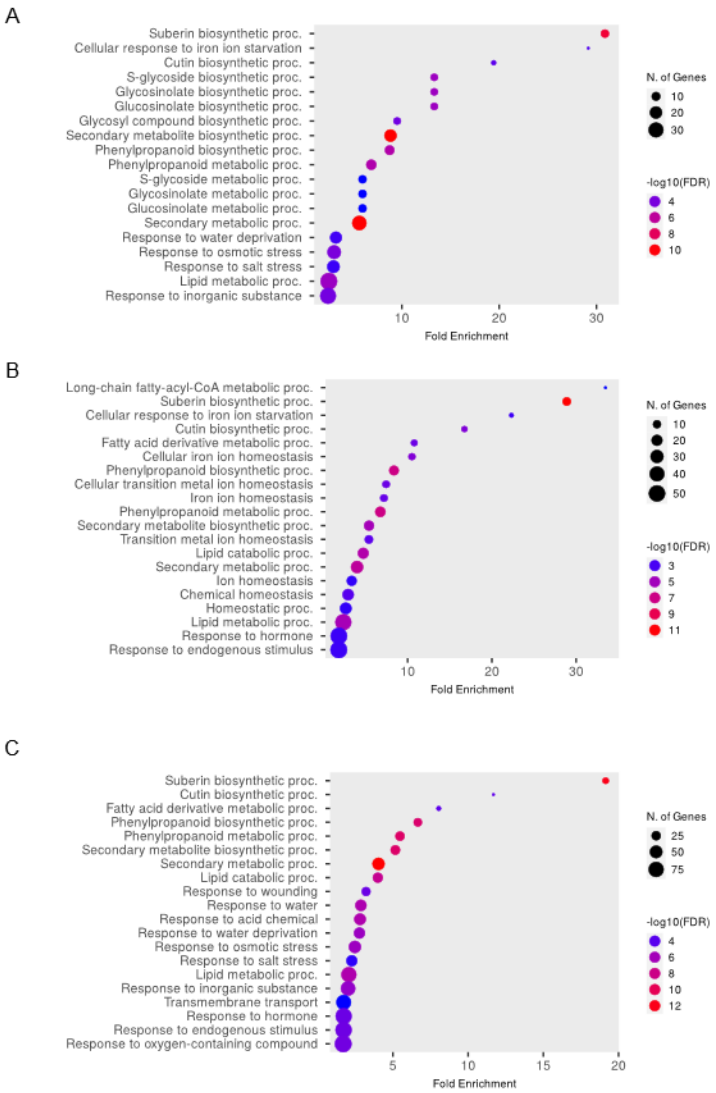

**Fig. S7 Gene Ontology terms enrichment in genes down-regulated in roots of N-degron pathway mutants.**

Differentially expressed gene lists were analysed with ShinyGO [17] using all genes identified in the RNA-seq data set as background. (A) *prt6-5*. (B) *big-2*. (C) *prt6-5 big-2*.

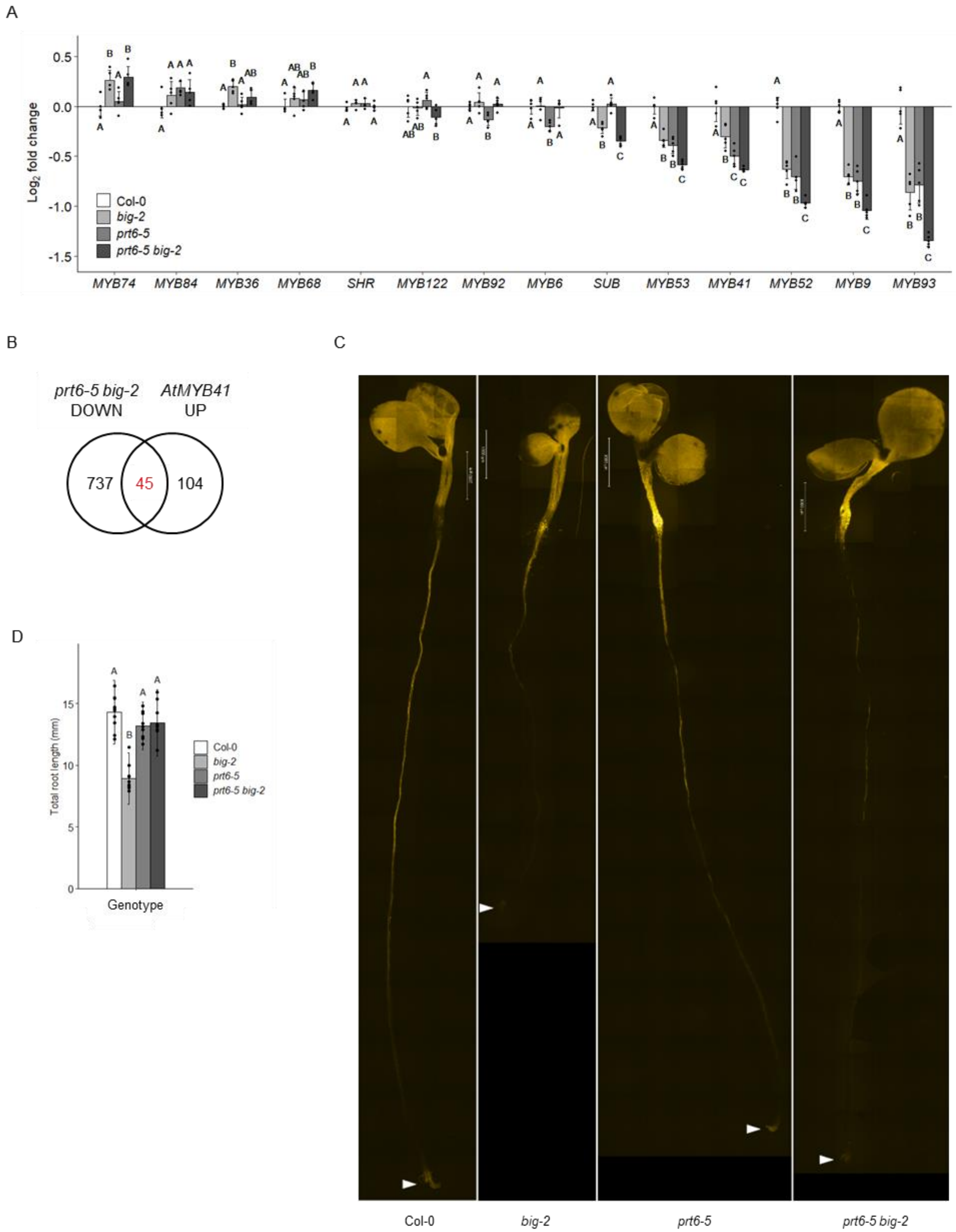

**Fig S8 suberin biosynthesis and deposition in N-degron pathway mutants.**

(A) Expression of transcription factors involved in suberin biosynthesis [18] relative to Col-0; data were re-plotted from the RNA-seq data set; different letters indicate significant differences between genotypes ($P < 0.05$). (B) Venn diagram showing comparison of genes down-regulated in *prt6-5 big-2* roots with genes up-regulated by ectopic expression of *MYB41* from [19]. (C) Representative composite micrographs showing Fluorol Yellow 088 staining of suberin in wild type and mutant roots (scale bar represents 1 mm). White arrowheads indicate positions of the root tips. (D) Root length of seedlings used for suberin staining; data represent means ± SD; different letters indicate significant differences between genotypes ($P < 0.05$).

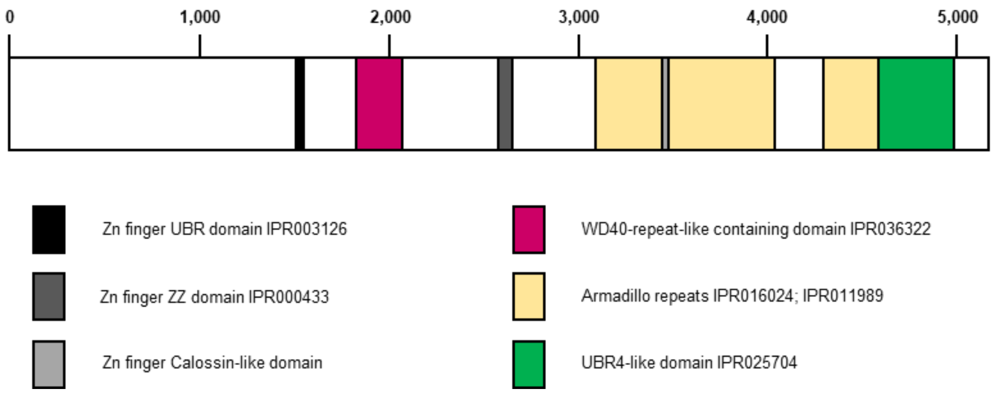

**Fig. S9 Cartoon of BIG showing positions of predicted protein domains.**

Domain positions as are indicated in [20] and TAIR. The scale shows number of amino acids.

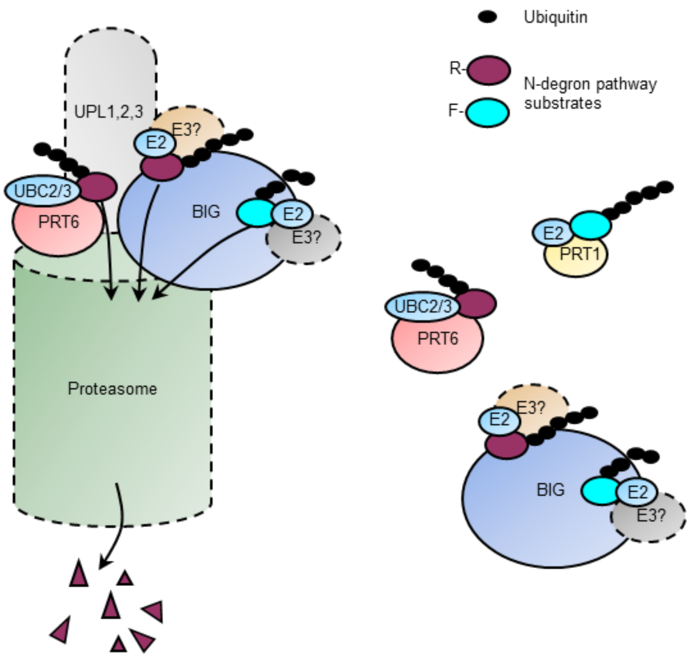

**Fig. S10 Speculative model for action of PRT6 and BIG**

The cartoon is based on information from the literature and data from the current study. A subset of the cellular pool of BIG and PRT6 is bound to the proteasome lid where the two proteins may interact [21, 22, 23, 24; Fig. 1F], but BIG and PRT6 likely also work without association with the proteasome. PRT6 is an E3 ligase that works together with the E2 conjugating enzymes, UBC2/3 to ubiquitinate protein substrates with basic (type 1) N-termini, targeting them for proteasomal degradation [1, 25]. BIG acts as a scaffold, recruiting one or more as-yet unidentified E3 (and E2) enzymes to ubiquitinate protein substrates with type 1 and type 2 N-termini, resulting in their degradation [26, 27, 28]; Fig. 1B,C]. Alternatively, BIG may have intrinsic E3 ligase activity. The PRT1 E3 ligase also mediates degradation of type 2 substrates [29]. Several reports indicate that the HECT-type ubiquitin protein ligases (UPLs) reside at the proteasome, where they increase the processivity of polyubiquitination in concert with multiple E3 ligases [30, 31]. BIG may also mediate degradation of proteins that are not substrates of the N-degron pathway and by analogy with UBR4 could participate in autophagy (not shown).

M/RRSGGGGGGGGGGGGRGGSGAWLLPVSLVKRKTTLAPNTQTASPRAL
ADSVNGKPIPNPLLGLDSTASKDNTVPLKLIALLANGEFHSGEQLGETLGMS
RAAINKHIQTLRDWGVDVFTVPGKGYSLPEPIPLLNAKQILGQLDGGSVAVLP
VVDSTNQYLLDRIGELKSGDACIAEYQQAGRGSRGRKWFSPFGANLYLSMF
WRLKRGPAAIGLGPVIGIVMAEALRKLGADKVRVKWPNDLYLQDRKLAGILV
ELAGITGDAAQIVIGAGINVAMRRVEESVVNQGWITLQEAGINLDRNTLAATLI
RELRAALELFEQEGLAPYLPRWEKLDNFINRPVKLIIGDKEIFGISRGIDKQGA
LLLEQDGVIKPWMGGEISLRSAEKLQLPPLERLTLDGGGGSGGGSMVSKGE
ELFTGVVPILVELDGDVNGHKFSVSGEGEGDATYGKLTLKFICTTGKLPVPW
PTLVTTFGYGLQCFARYPDHMKQHDFFKSAMPEGYVQERTIFFKDDGNYKT
RAEVKFEGDTLVNRIELKGIDFKEDGNILGHKLEYNYNSHNVYIMADKQKNGI
KVNFKIRHNIEDGSVQLADHYQQNTPIGDGPVLLPDNHYLSYQSALSKDPNE
KRDHMVLLEFVTAAGITLGMDELYK*

**Fig. S11 Amino acid sequence of R-Turbo and M-Turbo.**

Single letter amino acid code is used. The N-terminal residue revealed after cleavage *in planta* is indicated in red.

**Table S1. Genetic materials used in this study.**

| Line | Description | Source/reference |
|---|---|---|
| Col-0 | Wild type accession | NASC |
| *prt6-1* | T-DNA insertion mutant; SAIL_1278_H11 | 32 |
| *prt6-5* | T-DNA; SALK_0510_88 | 33 |
| *big-2* | T-DNA; SALK_045560 | 34 |
| *prt6-1 big-2* | | This study |
| *prt6-5 big-2* | | This study |
| *prt1-1* | EMS allele; Q111-STOP | 35; NASC accession, N119 |
| *ate1-2 ate 2-1 (ate1/2)* | T-DNA; SALK_023492<br>T-DNA; SALK_040788 | 32,33 |
| *ate1/2 big-2* | | This study |
| *rap2.12 rap2.2 rap2.3* | *rap2.12-1*, GABI-KAT line GK_503A1_11<br>*rap2.2-1*, SAIL184_G12<br>*rap2.3-1* allele; ref 36 | 6,9 |
| *prt6-1 rap2.12 rap2.2 rap2.3* | | 6 |
| *big-2 rap2.12 rap2.2 rap2.3* | | This study |
| *prt6-1 big-2 rap2.12 rap2.2 rap2.3* | | This study |
| *vrn2-5* | SALK_201153 | 4 |
| *prt6-1 vrn2-5* | | 4 |
| *big-2 vrn2-5* | | This study |
| *prt6-1 big-2 vrn2-5* | | This study |
| R-GUS Col-0 | | 1 |
| R-GUS *prt6-1* | | 1 |
| R-GUS *big-2* | | This study |
| R-GUS *prt6-1 big-2* | | This study |
| F-GUS Col-0 | | 1 |
| F-GUS *prt1-1* | | This study |
| F-GUS *big-2* | | This study |
| F-GUS *prt1-1 big-2* | | This study |
| M-GUS Col-0 | | 1 |
| M-GUS *prt1-1* | | 1 |
| M-GUS *prt6-1* | | This study |
| M-GUS *big-2* | | This study |
| M-GUS *prt6-1 big-2* | | This study |
| R-LUC Col-0 | | 3 |
| R-LUC *prt6-5* | | This study |
| R-LUC *big-2* | | This study |
| R-LUC *prt6-5 big-2* | | This study |
| p35S::RAP2.3-HA Col-0 | | 6 |
| p35S::RAP2.3-HA Col-0 *prt6-5* | | This study |
| p35S::RAP2.3-HA Col-0 *big-2* | | This study |
| p35S::RAP2.3-HA Col-0 *prt6-5 big-2* | | This study |
| p35S::HRE2-HA Col-0 | | 5 |
| p35S::HRE2-HA Col-0 *prt6-5* | | This study |
| p35S::HRE2-HA Col-0 *big-2* | | This study |
| p35S::HRE2-HA Col-0 *prt6-5 big-2* | | This study |

| | | |
|---|---|---|
| pVRN2::VRN2-GUS Col-0 | | This study |
| pVRN2::VRN2-GUS *prt6-1* | | 4 |
| pVRN2::VRN2-GUS *big-2* | | This study |
| pVRN2::VRN2-GUS *prt6-1 big-2* | | This study |
| pUBQ10::M-Turbo-NES-YFP (M-Turbo) Col-0 | | This study |
| pUBQ10::R-Turbo-NES-YFP (R-Turbo) Col-0 | | This study |

**Table S2. Primers used in this study**

| Purpose | Primer | Sequence (5'-3') |
|---|---|---|
| **Genotyping** | | |
| *big-2* | Big SALK_045560 LP1 | CGCATGGTGTCTTTGACCTAT |
| | Big SALK_045560 RP1 | CCCTGACATTGTTATGGTTGG |
| | LB1.3 | ATTTTGCCGATTTCGGAAC |
| *prt6-1* | AT120 | AAAATTGATCCTTTCCATGCC |
| | AT121 | CAACATAAGAATCTGCGGGAG |
| | LB2 SAIL | GCTTCCTATTATATCTTCCCAAATTACCAATACA |
| *prt6-5* | AT120 | AAAATTGATCCTTTCCATGCC |
| | AT121 | CAACATAAGAATCTGCGGGAG |
| | LB1.3 | ATTTTGCCGATTTCGGAAC |
| *prt1-1* (cleave product with *Mn* II) | N119_FP | CAGAGGAAGAGCAAGAACGAGAAT |
| | N119_RP | CCACCTTCTGTTTATCTACAC |
| *rap2.12* | rap2.12 GABI Fwd | CTCAGCTGTCTTGAACGTTCC |
| | rap2.12 GABI Rev | TGGCTACTCCTGAATGCAAAC |
| | GK8760 | GGGCTACACTGAATTGGTAGCTC |
| *rap2.2* | Sail_184 G12 RP | GAGCTAAGGACGGTAAGCTGG |
| | Sail_184 G12 LP | GAGAACATTTGTTCAATGCATG |
| | LB Sail | CTTCCTATTATATCTTCCCAAATTACCAATACA |
| *rap2.3* | AtEBP V2 LP | CAGTGAAGAAGGAGCAGGC |
| | AtEBP V2 RP | ATCCTATACACTCCACCGGG |
| | LB Sail | CTTCCTATTATATCTTCCCAAATTACCAATACA |
| *vrn2-5* | *vrn2-5*(SALK) LP | GTTTGTTCATCATGACACCCC |
| | *vrn2-5* (SALK) RP | TTTGAGTCACTGGGATGATCC |
| | LB1.3 (SALK) | ATTTTGCCGATTTCGGAAC |
| *ate1-2* | Ate1-2 F LG ate1-2 LP | TGAATGGCAGAATATTTTCCG |
| | Ate1-2 R LG ate1-2 RP | CAATTATGCATACGGAGTGGG |
| | LB1.3 | ATTTTGCCGATTTCGGAAC |
| *ate2-1* | Ate2-1 F At3 | GCGAAGCCGAGTGAGCAGACAGA |
| | Ate2-1 R At4-2 | CCACAAAGAGGAATCTTTTCTTCATCATCAT |
| | LB2 | CCAAACTGGAACAACACTCAACCCTATCTC |
| **qRT-PCR** (endogenous genes) | | |
| *ADH1* | ADH1_FW | GGTCTTGGTGCTGTTGGTTT |
| | ADH1_R | CTCAGCGATCACCTGTTGAA |
| *PGB1* | HB1_FW (PGB1) | GGCTCTTGTAGTGAAGTCTTGGA |
| | HB1_R (PGB1) | CTTCGTTGTTGGTGCAATCTCA |
| *HRE2* | HRE2_FW | TTGCTGCCATCAAAATCCGT |
| | HRE2_R | CCCCTGGTTTAGTATCGGCT |
| *RAP2.12* | RAP2.12_FW | ACTGAATGGGACGCTTCACTGG |

| | | |
|---|---|---|
| | RAP2.12_R | AGGGTTTGCACCATTGTCCTGAG |
| *RAP2.2* | RAP2.2_FW | CCTAGCGTCGTATCCCAGAA |
| | RAP2.2_R | CTCAGATGTGTTGGCTGCTG |
| *RAP2.3* | RAP2.3_FW | AACTCACGGCTGAGGAACTCTG |
| | RAP2.3_R | ACGTTAACTTGGTTGGTGGGATGG |
| *VRN2* | qPCR VRN2 P2 F | TCACTCTTCTGGTGTTGGAGA |
| | qPCR VRN2 P2 R | GATGGTGGCTGAGTCGACAA |
| **qRT-PCR** (transgene specific) | | |
| *HRE2* | HRE2_1654F | TATCGGAGGACCTGATGGCA |
| | HRE2_1772R | GGCCGCATTGGAGTCTTGAT |
| *RAP2.3* | RAP23_1939F | ATGAGTCCGTTTCCGAGGTG |
| | RAP23_2062R | TCGTAGGGGTAGGCGTAGTC |
| *GUS* | qPCR GUS F | TGTAGAAACCCCAACCCGTG |
| | qPCR GUS R | CTTGTAACGCGCTTTCCCAC |
| **qRT-PCR** (reference genes) | | |
| *ACT2* | ACT2 F | CCGCTCTTTCTTTCCAAGC |
| | ACT2 R | CCGGTACCATTGTCACACAC |
| *TUB4* | TUB4 F | GCTGAGAACAGCGATTGTCTT |
| | TUB4 R | AGTTCCCATTCCAGATCCAGT |
| AT5G18800 | qCTRL3_F | CCTTTTGGCACTTCTGGTG |
| | qCTRL3_R | GAAGTGTCTCGACAAAGGT |
| *UBQ10* | UBQ10F | GAAGTTCAATGTTTCGTTTCATGT |
| | UBQ10R | GGATTATACAAGGCCCCAAAA |
| **X-LUC assays** | | |
| R-LUC | KG59 | GCGGTCGGTAAAGTTGTTCCAT |
| | At327 | GAAGTGTTCGTCTTCGTCCC |
| AT2G28390 (reference gene) | MON1_up | AACTCTATGCAGCATTTGATCCACT |
| | MON1_lo | TGATTGCATATCTTTATCGCCATC |

**Dataset S1 TurboID proteomics data.**

**Dataset S2 RNA-seq data.**
